# Determination of both global and local protein stability in complex environments using X-ray hydroxyl radical footprinting

**DOI:** 10.64898/2026.11.21.768341

**Authors:** Shawn M. Costello, Darren N. Kahan, Mark Petersen, Sayan Gupta, Corie Y. Ralston, Susan Marqusee

## Abstract

Protein thermodynamic stability is a major determinant of cellular function, yet traditional methods for determining protein stability require large quantities of pure protein under idealized conditions. Here, we overcome this challenge by combining X-ray hydroxyl radical footprinting monitored by mass spectrometry (XF-MS) with chemical denaturation, to accurately measure protein thermodynamic stability. We demonstrate that XF-MS quantitatively reproduces known stability values using ∼500-fold less protein than traditional spectroscopic approaches such as circular dichroism. Detection with mass spectrometry extends these measurements to heterogeneous samples demonstrated by measuring the stability of individual proteins in complex mixtures, including in cell lysate. Furthermore, by monitoring XF-MS at the level of individual peptides, we can monitor local stability and detect deviations from the typical assumed two-state unfolding behavior. This local resolution allows us to detect distinct cooperative units and assign thermodynamic values to individual structural domains within a multi-domain protein. In sum, XF-MS extends quantitative stability measurements to a broader range of proteins, environments, and levels of structural detail than previously accessible with conventional approaches.

**Significance Statement:** A protein’s thermodynamic stability, or how strongly it favors its folded over its unfolded state, determines the fraction of functional molecules and the fraction prone to misfolding, aggregation, and degradation. Rigorous quantification of stability typically requires milligrams of purified protein studied in isolation, restricting studies to a few proteins under artificial conditions. We show that X-ray hydroxyl radical footprinting, which irreversibly labels solvent-exposed side chains within microseconds, combined with chemical denaturation and mass spectrometry, accurately measures stability using ∼500-fold less protein. Because mass spectrometry separates each protein and peptide, the approach monitors individual proteins within mixtures and cell lysate and resolves which regions unfold together. This opens thermodynamic studies to larger proteins and more realistic environments.

## Introduction

Proteins are large and complex macromolecules that perform a wide range of essential biological functions. Each protein’s amino acid sequence, in the context of its environment, encodes its fold and function (1–3). Advances in AI have created a revolution in our ability to predict a protein’s fold given only its amino acid sequence with accuracy comparable to experiment (4–9). To accomplish their biological function, however, proteins need to access more than a single folded structure; proteins are dynamic, sampling alternative conformations with defined populations and rates of interconversion. Together, these conformations and their dynamics are referred to as a protein’s conformational energy landscape (10). Features of this energy landscape are critical determinants of enzyme catalysis, allostery, signaling, misfolding, and turnover, among many other functions (10, 11). Therefore, predicting both fold and function from protein sequence and environment requires a deeper understanding of the energy landscape. The most basic description of a protein’s energy landscape is its global stability 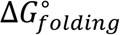, defined as the difference in free energy between the native, folded state and the largely unstructured, unfolded state. While a simple parameter, changes in global stability can have effects that range from undetectable to pathological.

Global stability can be quantified by the ratio of folded to unfolded molecules of a given protein at equilibrium 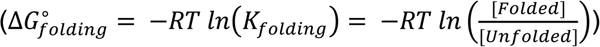 and can vary by orders of magnitude depending on the protein. For stable proteins, this ratio can be greater than 10^8^ 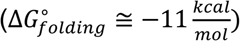, whereas for unstable proteins, it can be 1 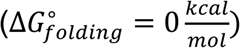 or lower (12–15). Small changes in environment or sequence (e.g., single-site variants) can dramatically alter protein stability, which can have important consequences for biology, medicine, and industrial use (13, 16). The populations of both states—the native and unfolded—are important determinants of protein function. Native states allow for functional binding or catalysis, and non-native unfolded conformations can misfold, aggregate, or be degraded by proteases. Therefore, even when destabilization has minor effects on native-state structure, catalysis, or binding, it can still greatly affect function by modulating steady-state abundance of active molecules and by promoting toxic aggregation (16–18).

Measuring the relative population of unfolded and folded molecules is an experimental challenge, in part because globular proteins frequently have 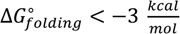 (which would require detecting the <1 in 100 molecules that are unfolded). Traditional approaches rely on perturbation, using chemical denaturants or temperature, and monitoring the populations throughout the unfolding transition (where the ratio of folded:unfolded molecules is between 10:1 and 1:10), and then extrapolating to native conditions (the absence of perturbant) (19). In principle, any technique capable of measuring the ratio of folded:unfolded proteins can be used to determine protein stability, as long as the measurement technique itself does not perturb this equilibrium population (20). The most common approaches rely on spectroscopic probes of structure, such as circular dichroism (CD) or intrinsic fluorescence, and require significant amounts (usually milligrams) of pure protein. Thus, quantitative studies of protein stability are typically limited to abundant proteins that can be easily generated, usually via overexpression in *E. coli*, and purified. Because all components in the sample contribute to the measured signal, such stability measurements cannot be made in the presence of other biomolecules, limiting most measurements of protein stability to simple model systems under idealized, rather than realistic, physiological conditions.

Analysis of these data also requires simplifying assumptions. The most common is the two-state assumption: that the protein only populates two states, native and unfolded (19). As protein folding is a highly cooperative process, this is often a valid assumption for single-domain proteins. However, for larger proteins and proteins with multiple domains, partially folded states are often stable enough to be populated in the unfolding transition, requiring more complex models and the need to assess local or regional stability rather than the simple parameter of global stability (21–23).

The most promising approaches to measure stability at a local level rely on selective labeling, or modification, at individual sites, such as hydrogen-deuterium exchange (HDX), pulsed proteolysis, and oxidative footprinting (24–27). The local nature of these probes can help distinguish between two-state and multi-state unfolding. Detection of these modifications using mass spectrometry or polyacrylamide gel electrophoresis (PAGE) can overcome sample quantity and purity limitations. Each of these methods, however, brings its own restrictions. For example, HDX is less suitable for monitoring proteins in complex conditions - the reversible nature of the deuterium label limits the ability to carry out any biochemistry or separation processes before the detection phase. For pulse proteolysis, the native state of the protein cannot have any protease-susceptible flexible loops, and for oxidation-based methods, the modification itself may perturb the folding equilibrium. Finally, in any pulsed-labeling methods, the protein must not be able to interconvert between conformations on the timescale of the labeling pulse (typically seconds to minutes), which limits their applicability.

To overcome these challenges, we turned to another selective modification approach, X-ray hydroxyl radical footprinting monitored by mass spectrometry (XF-MS). XF-MS is a powerful tool that uses short exposures (tens to hundreds of microseconds) of synchrotron-generated X-ray beams to produce hydroxyl radicals from water molecules in solution (28–31). The resulting radicals rapidly and irreversibly modify nearby amino acid side chains. The extent of hydroxyl radical labeling (oxidation) on a protein is known to be a direct result of both the sidechain (the chemistry of the reactive group) and the solvent-accessible surface area of that side chain (27, 31). This dependence on solvent accessibility has made hydroxyl radical footprinting an effective structural probe for protein conformations and protein-protein interactions (28, 31–36). This feature of XF-MS makes it well-suited for measuring the ratio of folded:unfolded proteins, as it differentially labels states based on their exposure of buried side chains.

XF-MS has several advantages over other approaches for monitoring protein folding and unfolding. Because the oxidation is monitored by MS, the measurements require minimal sample (sub-microgram) and can be carried out without extensive purification. Since the oxidation is irreversible, samples in biological conditions less suitable for MS detection can be further purified before analysis. Moreover, because labeling times are shorter than the timescales of most folding and unfolding reactions, XF-MS represents a snapshot of the relative populations under the conditions of the labeling pulse. Lastly, as XF-MS monitors structure at the level of individual peptides, it provides information about the local or regional stability within a protein.

Here, we demonstrate that XF-MS can be used to accurately measure the thermodynamic stability of a protein in purified, idealized conditions, and, in addition, can be used to measure the stability of the protein directly from *E. coli* lysate, in mixtures with another protein, and locally at the level of individual peptides. We then extend this study to a multi-domain non-two-state protein, demonstrating the ability to monitor multi-state unfolding and structurally assign measured thermodynamic stability to specific domains within larger proteins. XF-MS expands the ability to measure conformational stability to important biological systems and contexts and should be sensitive enough to structurally probe unstable or even intrinsically disordered proteins.

## Results

### XF-MS differentially labels the folded and unfolded state of a protein

In order to serve as a measure of protein stability, XF-MS must differentially label proteins in the folded and unfolded state. Since the short-lived radicals generated from solvent react with nearby side chains, solvent accessibility is a major determinant of the oxidation reaction rate (27, 37). After cysteine, large aromatic and hydrophobic side chains, such as tryptophan, phenylalanine, methionine, and tyrosine, react most rapidly with the hydroxyl radicals (31). These same residues are usually buried in the cores of folded proteins; therefore, it stands to reason that the folded and unfolded state would be differentially modified during a short pulse of hydroxyl radical labeling.

To develop and test the ability of XF-MS to monitor protein denaturation, we used the well-characterized single-domain protein, *E. coli* RNase H (ecRNH*, where the * indicates all cysteines have been replaced with alanine) (38). ecRNH* was pre-equilibrated in differing concentrations of urea, and X-ray exposure was performed by flowing each sample through a capillary tube perpendicular to a synchrotron beam such that the duration of exposure (pulse length) was 500 or 1000 microseconds. The degree of protein oxidation in each sample was then measured by MS. In increasing concentrations of urea (increasing the population of the unfolded state), the observed mass increases, consistent with increased oxidation of exposed side chains (Fig. 1A and *SI Appendix* Fig. S1). Thus, as expected, hydroxyl radicals generated by XF-MS differentially react with the unfolded and folded conformations.

**Figure 1.**
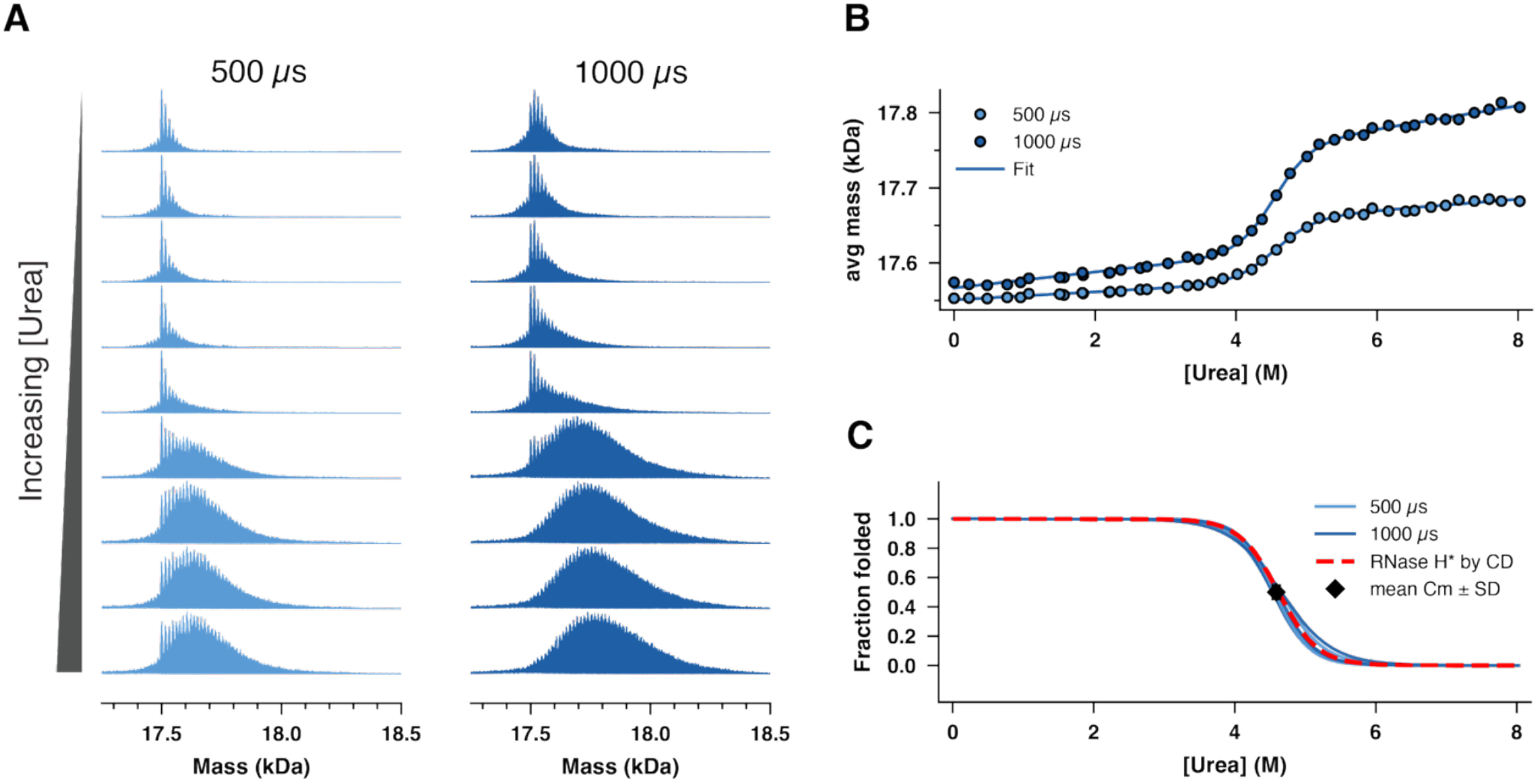
XF-MS denaturant melts of ecRNH* quantitatively reproduce the global stability measured by traditional probes. (A) Deconvoluted mass spectra for ecRNH* (50 mM KCl, 20 mM NaOAc, pH 5.5, room temperature) for a representative subset of urea concentrations for two different exposure times, 500 and 1000 µs. (B) Average mass for deconvoluted mass spectra as a function of denaturant for a single representative denaturant melt at two different exposure times, 500 and 1000 µs. (C) Overlay of triplicate denaturant melts at two exposure times, normalized to fraction folded. The expected fraction folded determined by circular dichroism is shown as a dotted red line.

### Chemically induced denaturation monitored by XF-MS quantitatively reproduces known thermodynamic stability using >500-fold less sample than traditional approaches

The equilibrium stability of ecRNH* has been measured with several different techniques (25, 38–41). To compare to these previous studies, we carried out XF-MS as a function of urea. When these mass spectra at each denaturant condition are quantified by determining the average of the mass distributions, a clear cooperative transition is observed (Fig. 1B and *SI Appendix* Fig. S2). These data can be fit using a traditional two-state linear-extrapolation model (LEM, see methods) (19) to obtain free energies of folding 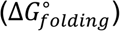 and m-values (dependence of 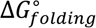 on urea) that agree with the stability determined by more traditional methods such as CD (Fig. 1C, Table 1) (41). Thus, XF-MS accurately determines global stability in a simple two-state model system.

**Table 1.**
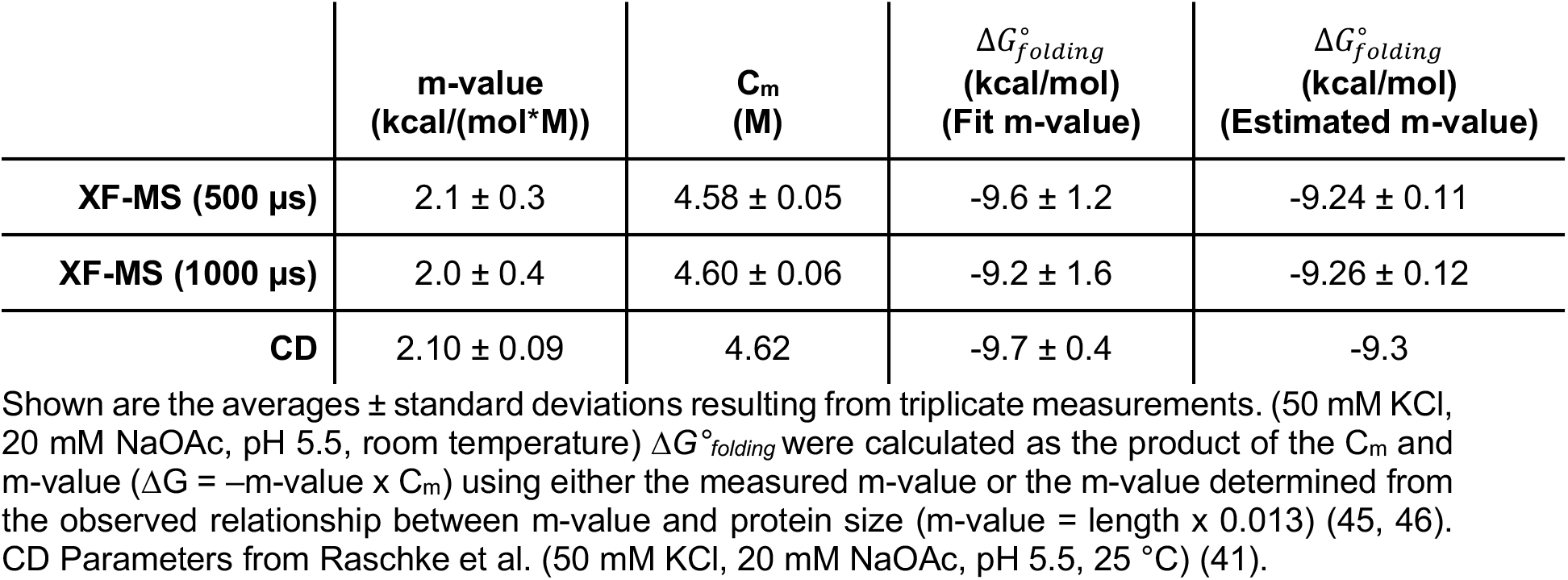
Thermodynamic parameters from ecRNH* XF-MS denaturation.

As with all measurements of protein stability, a concern is that the measurement itself may perturb the equilibrium. Thus, it is essential to make sure that oxidation is not inducing unfolding over the timescale of the labeling reaction, allowing for further oxidation, which would result in a lower apparent stability. The agreement between the melts obtained from two different pulse lengths (500 and 1000 microseconds, Fig. 1C, Table 1) supports the assumption that modifications are not inducing unfolding on the timescale of the experiment. Additionally, the agreement between XF-MS and CD further supports the validity of XF-MS as a method for monitoring denaturation. Based on a set of previously well-characterized two-state folding proteins, we expect that most proteins will not significantly re-equilibrate during the one-millisecond pulse used in XF-MS (12). Remarkably, this experiment used only 0.1 micrograms of protein per denaturant condition, compared to traditional optical methods which typically use approximately 50 micrograms per denaturant condition. Thus, XF-MS requires at least 500-fold less sample than CD and other spectroscopic methods.

### XF-MS can monitor the chemically induced denaturation of multiple proteins simultaneously

A major limitation of traditional spectroscopic approaches is the need for highly pure single-component samples. With a mixture of proteins, or contaminants, each will contribute to the observed spectroscopic signal. When samples contain two or more proteins, the observed results will produce either multiple transitions (which can be difficult to assign to individual components) or if the transitions for each protein occur at similar denaturant concentrations or temperatures, a single broadened transition, which (when erroneously analyzed as a single protein) returns thermodynamic values that are inconsistent with any of the proteins in solution. LC/MS-based approaches are ideally suited to handle this heterogeneity. Thus, XF-MS should be amenable to any protein mixture if the proteins of interest (or peptides resulting from the digestion of those proteins) are separable by LC/MS. To demonstrate this functionality, we monitored the equilibrium chemically induced denaturation of a mixture of two proteins, ecRNH* and T4 lysozyme (T4L*, where * represents a cysteine-free construct) by both XF-MS and CD.

When monitored by CD, the observed transition of this 1:1 mixture looks like a single, albeit broad, cooperative transition that is adequately fit by a single-protein two-state analysis (*SI Appendix*, Fig S3). This transition does not reflect the behavior of either component in the system, nor does it reflect any linear combination or average of the two. When fit using a traditional LEM, the resulting stability is a nonsensical 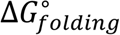 (-7.0 kcal/mol), lower than either of the 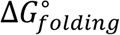 for ecRNH* or T4L* (Table 2).

**Table 2.**
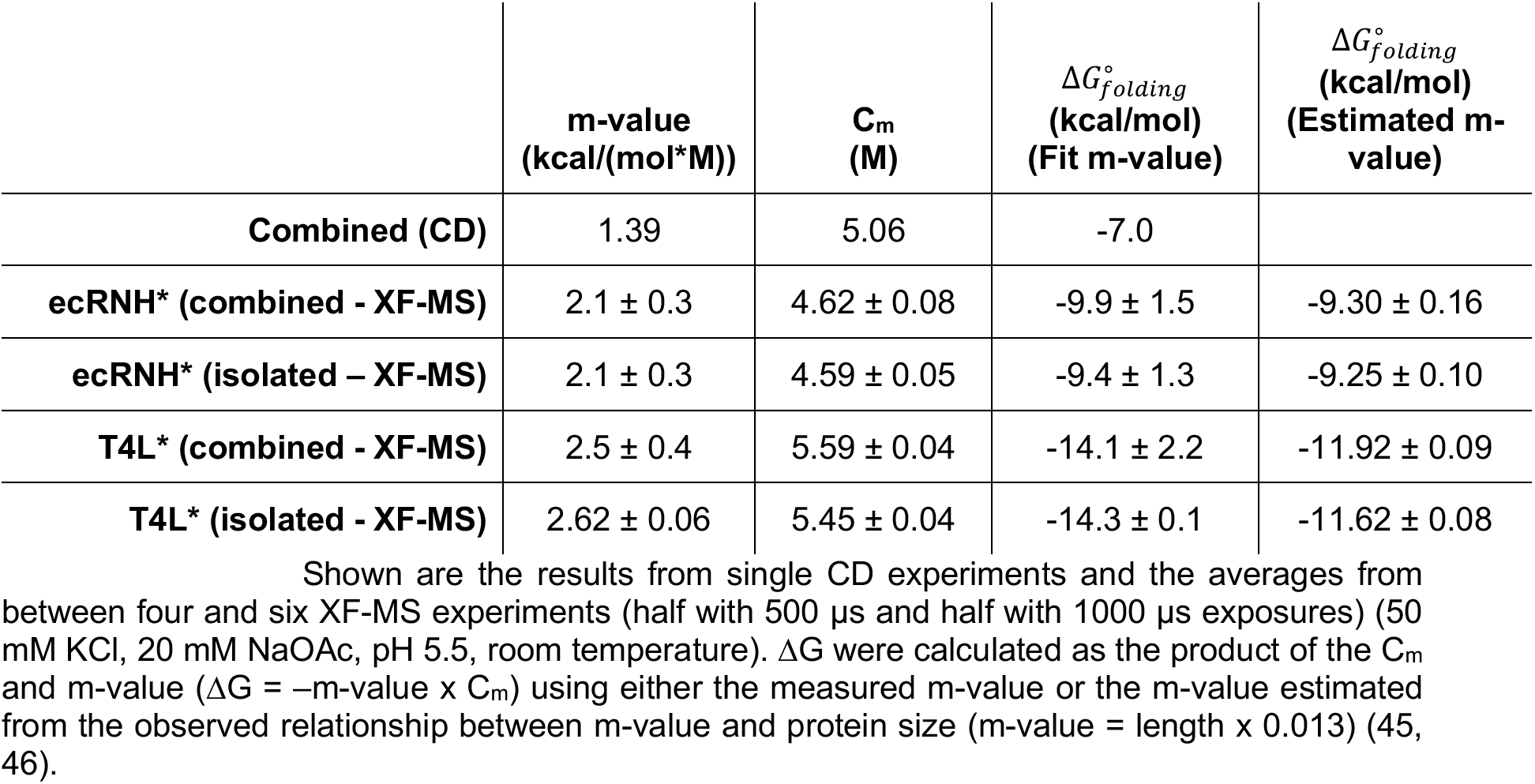
Thermodynamic parameters from the isolated and combined denaturation of ecRNH* and T4L*.

|  | **m-value<br>(kcal/(mol\*M))** | **C<sub>m</sub><br>(M)** | Δ*G*<sup>°</sup><sub>*folding*</sub><br>**(kcal/mol)<br>(Fit m-value)** | Δ*G*<sup>°</sup><sub>*folding*</sub><br>**(kcal/mol)<br>(Estimated m-value)** |
|---|---|---|---|---|
| **Combined (CD)** | 1.39 | 5.06 | -7.0 |  |
| **ecRNH\* (combined - XF-MS)** | 2.1 ± 0.3 | 4.62 ± 0.08 | -9.9 ± 1.5 | -9.30 ± 0.16 |
| **ecRNH\* (isolated – XF-MS)** | 2.1 ± 0.3 | 4.59 ± 0.05 | -9.4 ± 1.3 | -9.25 ± 0.10 |
| **T4L\* (combined - XF-MS)** | 2.5 ± 0.4 | 5.59 ± 0.04 | -14.1 ± 2.2 | -11.92 ± 0.09 |
| **T4L\* (isolated - XF-MS)** | 2.62 ± 0.06 | 5.45 ± 0.04 | -14.3 ± 0.1 | -11.62 ± 0.08 |

Shown are the results from single CD experiments and the averages from between four and six XF-MS experiments (half with 500 µs and half with 1000 µs exposures) (50 mM KCl, 20 mM NaOAc, pH 5.5, room temperature). ΔG were calculated as the product of the C<sub>m</sub> and m-value (ΔG = –m-value x C<sub>m</sub>) using either the measured m-value or the m-value estimated from the observed relationship between m-value and protein size (m-value = length x 0.013) (45, 46).

Because ecRNH* and T4L* have non-overlapping mass distributions (Fig 2A and *SI Appendix*, Fig. S4), XF-MS reveals two distinct transitions and allows independent determination of the stability of each protein in the mixture. (Fig. 2B and *SI Appendix*, Fig. S5). The resulting thermodynamic values are consistent with the stabilities for both ecRNH* and T4L* measured in isolation (Table 2), indicating, as expected, that the stability of each protein is not affected by the presence of the other. Thus, XF-MS provides a unique method for quantitatively determining the stability of individual components within a mixture, providing ease in studying complex mixtures and the ability to identify and quantify any interactions.

**Figure 2.**
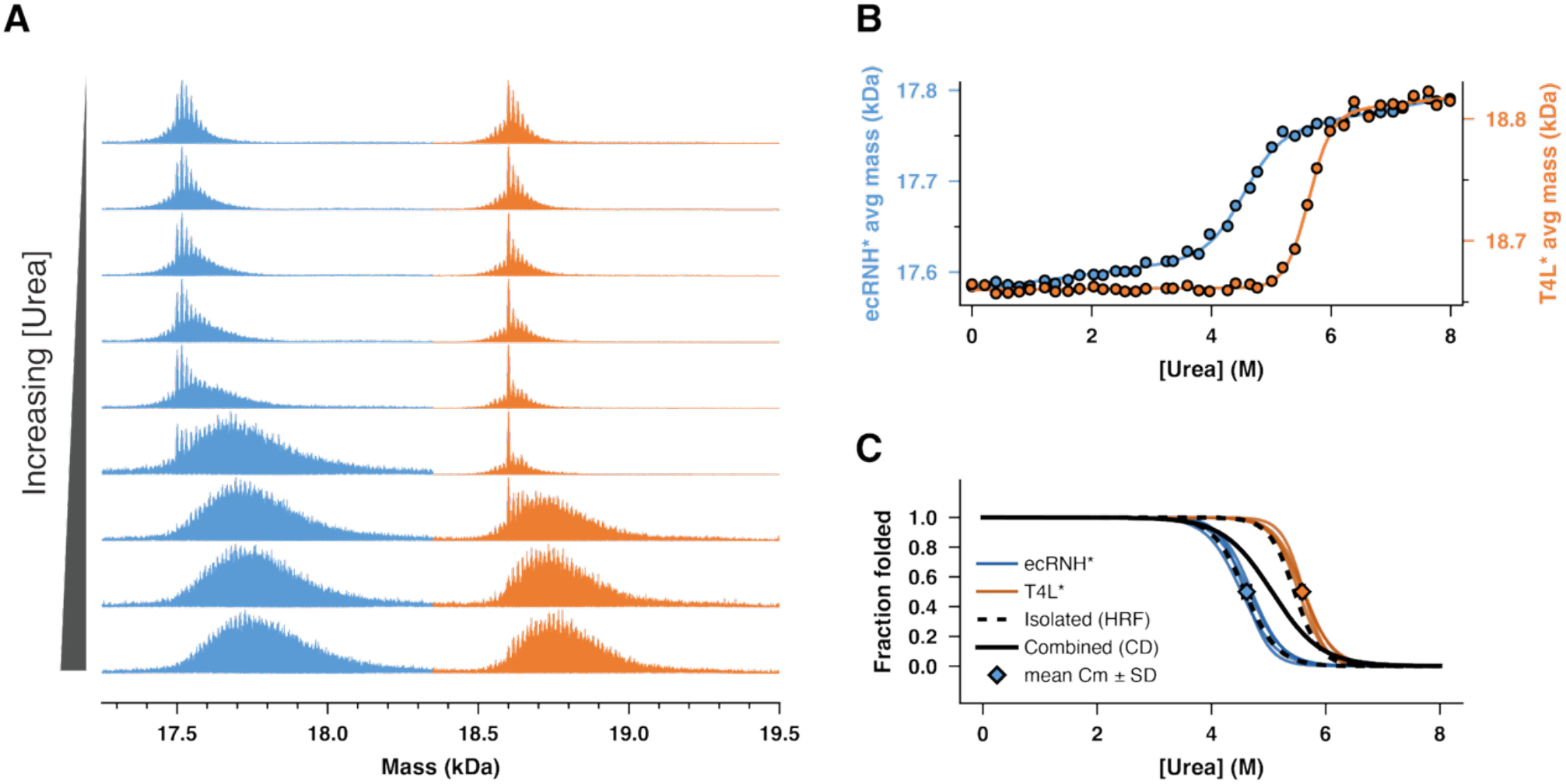
XF-MS separately resolves the unfolding of ecRNH* and T4L* within a mixture. (A) Deconvoluted mass spectra for a mixture of ecRNH* (blue) and T4L* (orange) (50 mM KCl, 20 mM NaOAc, pH 5.5, room temperature) for a representative subset of varying concentrations of denaturant after a 1000 µs x-ray exposure. Blue and orange data are normalized independently for ease of visualization. (B) Average mass of ecRNH* (blue) and T4L* (orange) deconvoluted mass spectra as a function of denaturant for a single representative denaturant melt collected at a 1000 µs exposure times. (C) Overlay of denaturant melts at two different exposure times (500 µs and 1000 µs) normalized to fraction folded. Melts obtained by XF-MS are shown for ecRNH* (blue) and T4L*(orange). A melt of an equimolar mixture of ecRNH* and T4L* obtained by CD is shown as a black solid line. Expected fraction folded for isolated ecRNH* and T4L* based on denaturant melts monitored by CD are shown as dotted lines.

### XF-MS can monitor the chemically induced denaturation of proteins directly in cell lysate

Because XF-MS uses mass spectrometry and monitors an irreversible modification, the approach is amenable to proteins in lysates or other complex biological settings. In addition to bypassing purification steps, this allows for stability measurements of proteins in biological settings that are not amenable to other methods. Should further purification be required, the irreversible nature of the modification allows for manipulation or enrichment of the sample after labeling, but before quantification. To demonstrate the feasibility of this approach in complex conditions, we performed a denaturant melt of ecRNH* in clarified *E. coli* lysate. The soluble components of *E. coli* lysate were isolated, buffer exchanged into assay buffer (50 mM KCl, 20 mM NaOAc, pH 5.5), and diluted into varying concentrations of denaturant with 1 µM ecRNH* added (see methods) and XF-MS was performed.

The resulting spectra show a considerably lower extent of modification for ecRNH* (Fig. 3A and *SI Appendix*, Fig. S6) compared to the same exposure and denaturant conditions in the absence of lysate (Fig.1A). This is apparent when comparing the resulting denaturant melts (Fig. 3B and *SI Appendix*, Fig. S7), where the change in average mass is approximately five-fold lower than what was observed for isolated ecRNH*. This is expected, as the additional macromolecules present in the lysate will react with the generated hydroxyl radicals (i.e. quenching) (42). This reduced steady-state concentration of radicals can be compensated by longer exposure times or improvements in beamline infrastructure, including recently developed container-free exposure approaches (43, 44). Even with this decrease in signal, the denaturant melts in lysate are quantifiable and can be compared to isolated ecRNH* (Fig. 3C). The C_m_ measured in lysate is slightly lower than the C_m_ measured in isolation (Welch p = 0.047, n = 3), indicating interactions between ecRNH* and other components in the lysate may result in minor destabilization (Table 3). However, the large error in the m-value, could indicate a change in the cooperativity of ecRNH* folding in lysate. These results demonstrate the ability of XF-MS to monitor protein unfolding in lysate, increasing throughput by reducing the time and resources required for more extensive purification, and suggests the possibility of performing more detailed thermodynamic studies of protein folding in complex biological environments previously thought to be intractable.

**Figure 3.**
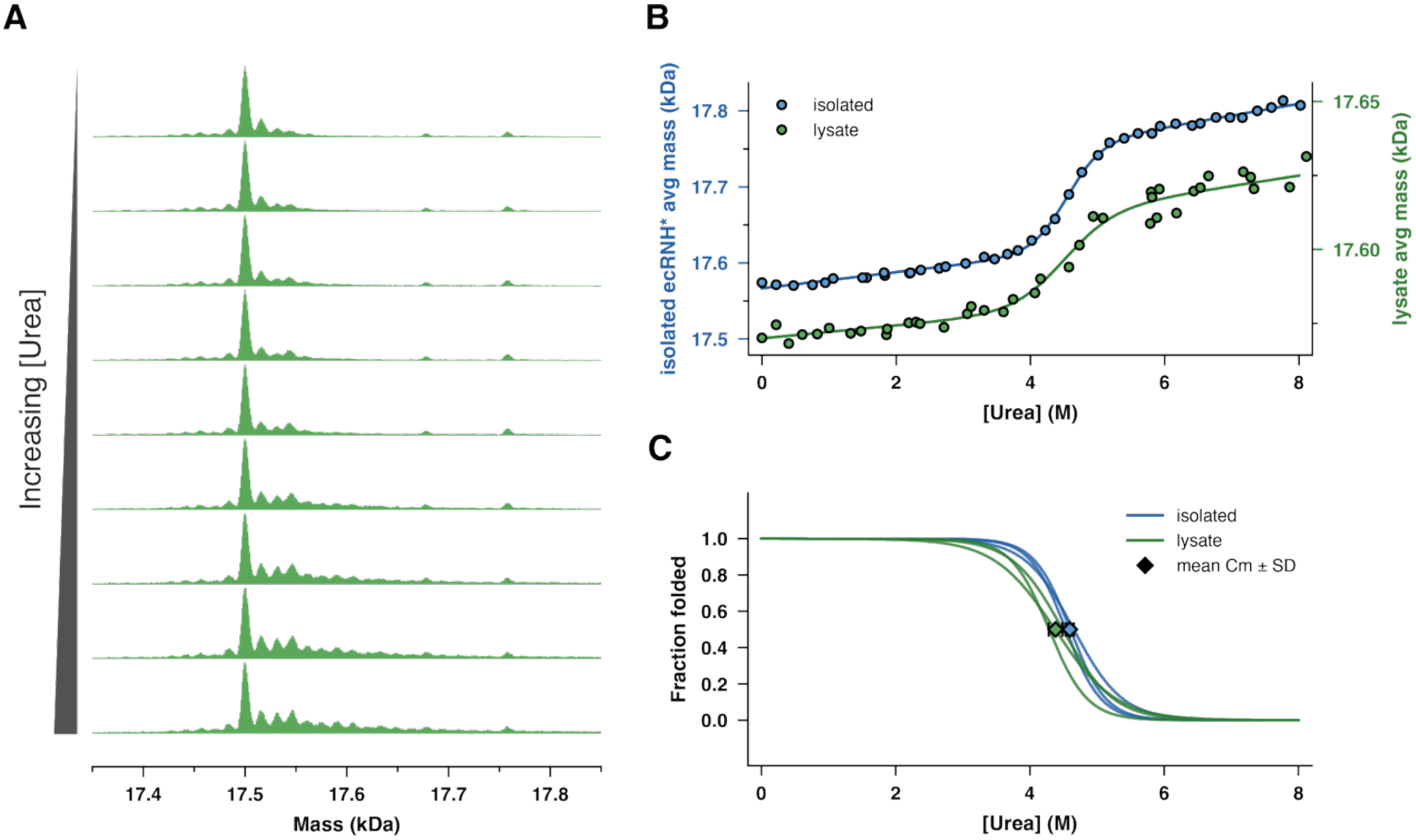
XF-MS measures the denaturant melt of ecRNH* directly in *E. coli* lysate. (A) Deconvoluted mass spectra for purified ecRNH* diluted into *E. coli* lysate in a representative subset of varying concentrations of denaturant for 1000 µs exposure time. (B) Average mass of isolated ecRNH* (blue) and ecRNH* in *E. coli* lysate (green) deconvoluted mass spectra as a function of denaturant for a single representative denaturant melt. Data are shown offset on separate y-axes for ease of visualization. (C) Overlay of triplicate denaturant melts normalized to fraction folded. Melts obtained by XF-MS are shown for ecRNH* (blue) and ecRNH* in *E. coli* lysate (green).

**Table 3.**
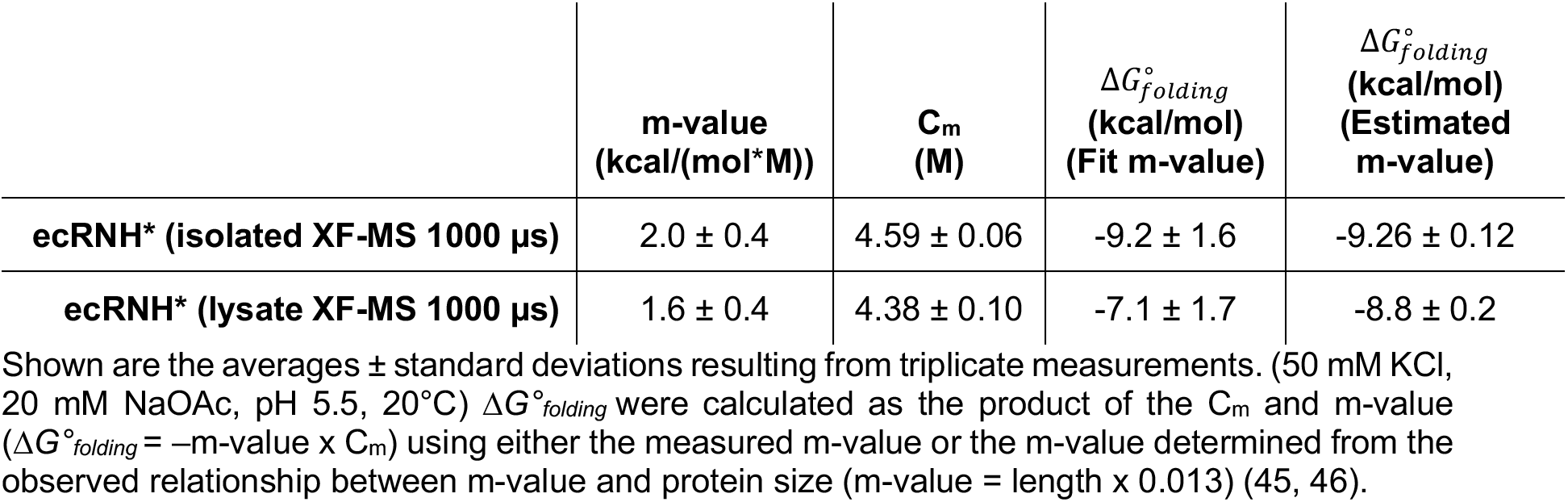
Thermodynamic parameters from ecRNH* XF-MS denaturation in the presence and absence of clarified cell lysate.

|  | **m-value<br>(kcal/(mol\*M))** | **C<sub>m</sub><br>(M)** | ΔG°<sub>folding</sub><br>**(kcal/mol)<br>(Fit m-value)** | ΔG°<sub>folding</sub><br>**(kcal/mol)<br>(Estimated m-value)** |
|---|---|---|---|---|
| **ecRNH\* (isolated XF-MS 1000 µs)** | 2.0 ± 0.4 | 4.59 ± 0.06 | -9.2 ± 1.6 | -9.26 ± 0.12 |
| **ecRNH\* (lysate XF-MS 1000 µs)** | 1.6 ± 0.4 | 4.38 ± 0.10 | -7.1 ± 1.7 | -8.8 ± 0.2 |

Shown are the averages ± standard deviations resulting from triplicate measurements. (50 mM KCl, 20 mM NaOAc, pH 5.5, 20°C) Δ*G*°<sub>*folding*</sub> were calculated as the product of the C<sub>m</sub> and m-value (Δ*G*°<sub>*folding*</sub> = –m-value x C<sub>m</sub>) using either the measured m-value or the m-value determined from the observed relationship between m-value and protein size (m-value = length x 0.013) (45, 46).

### XF-MS can monitor the chemically induced denaturation of proteins locally at the peptide level

As demonstrated above, using XF and monitoring the chemically induced denaturation by intact MS returns accurate thermodynamic parameters. However, this approach relies on the ability to obtain mass spectra of full-length protein and provides only global thermodynamic parameters. For some proteins, especially very large proteins, following the intact protein mass spectra can be technically challenging, and following individual peptides may be a simpler approach. Importantly, while the above intact protein analysis yields measurements of global stability, they may obscure subtle differences throughout the molecule, such as the presence of partially folded intermediates. Thus, we performed the same chemical denaturation XF-MS experiment but assessed modifications at a local level; after exposure to X-rays, we digested the sample (using trypsin) and measured the oxidation susceptibility at the level of individual peptides (see Methods). Of the eleven tryptic peptides detected for ecRNH*, nine showed cooperative transitions consistent with unfolding (Fig. 4A and *SI Appendix*, Fig. S8). The two peptides without a cooperative transition are located in regions that are solvent exposed in the folded state and are therefore not expected to have a change in side-chain solvent accessibility, or susceptibility to oxidation, upon unfolding. These peptides with transitions provide local measurements of folding stability at the corresponding regions in the protein structure.

**Figure 4.**
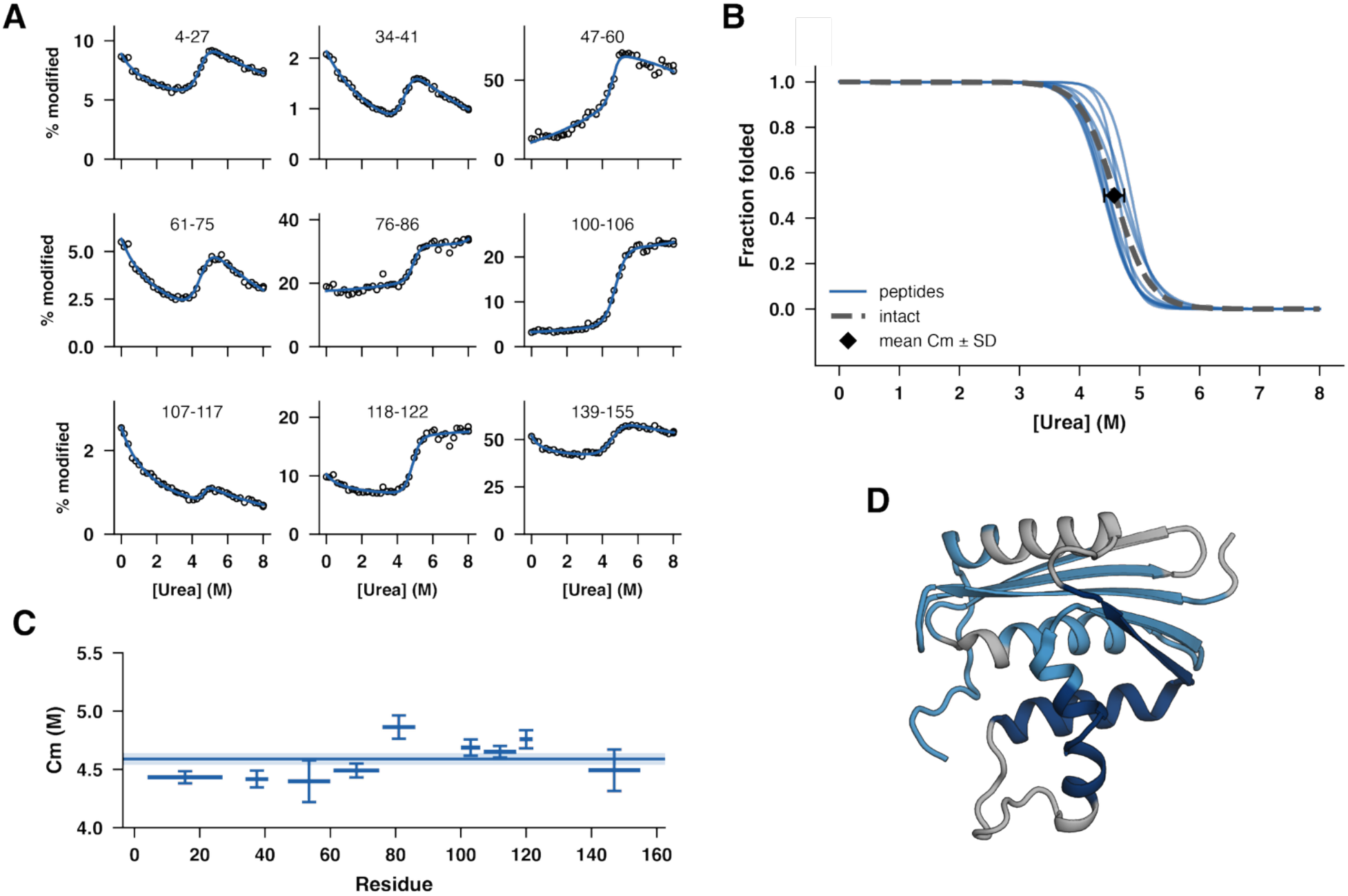
Peptide-level XF-MS provides multiple independent probes of ecRNH* folding. (A) Percent modified vs. denaturant concentration for nine tryptic peptides from ecRNH* for a single representative denaturant melt. Residue numbers are indicated in each panel. Blue lines represent fits to a modified LEM (see methods). (B) Overlay of denaturant melts for all tryptic peptides with observable transitions, using the average C_m_ and m-value determined from triplicate measurements for each peptide, normalized to fraction folded. The expected fraction folded based on intact melts shown in Figure 1 is shown as a blue line. (C) Concentration midpoints (C_m_) for each tryptic peptide. Sequence position for each peptide is represented by the x-axis. Error bars represent the standard deviation determined from triplicate measurements. Solid lines indicate values from intact melts (figure 1), with shaded regions representing the standard deviation determined from triplicate measurements. (D) Structure of ecRNH* (PDB: 2RN2) with peptides colored by C_m_ (C_m_ < global C_m_ as light blue, C_m_ > global C_m_ as dark blue, peptides with no cooperative transitions, or regions without peptide coverage, as gray)

Analysis of these peptides proved to be more complex than for the intact protein. Although each has a cooperative transition, the plots of peptide modification as a function of denaturant showed peptide-specific non-linear baselines (presumably due to non-linear denaturant quenching effects). Therefore, instead of using a traditional six-parameter LEM with linear baselines, we used a modified seven-parameter LEM, where we allow for a non-linear denaturant dependent baseline (see Methods). The resulting denaturation profiles overlay well with the global melt obtained by intact XF-MS (Fig. 4B). The C_m_ determined for each peptide showed higher variation than the C_m_ determined for the intact protein, and each fit had poorly determined m-values (*SI Appendix*, Fig. S9). This is not unexpected since this modified model increases the total number of fit parameters from six to seven, increasing uncertainty. Nonetheless, using this analysis, all peptides have denaturant concentration midpoints (C_m_) within 0.3 M of the global intact melt (Fig. 4C, Table 4).

**Table 4.**
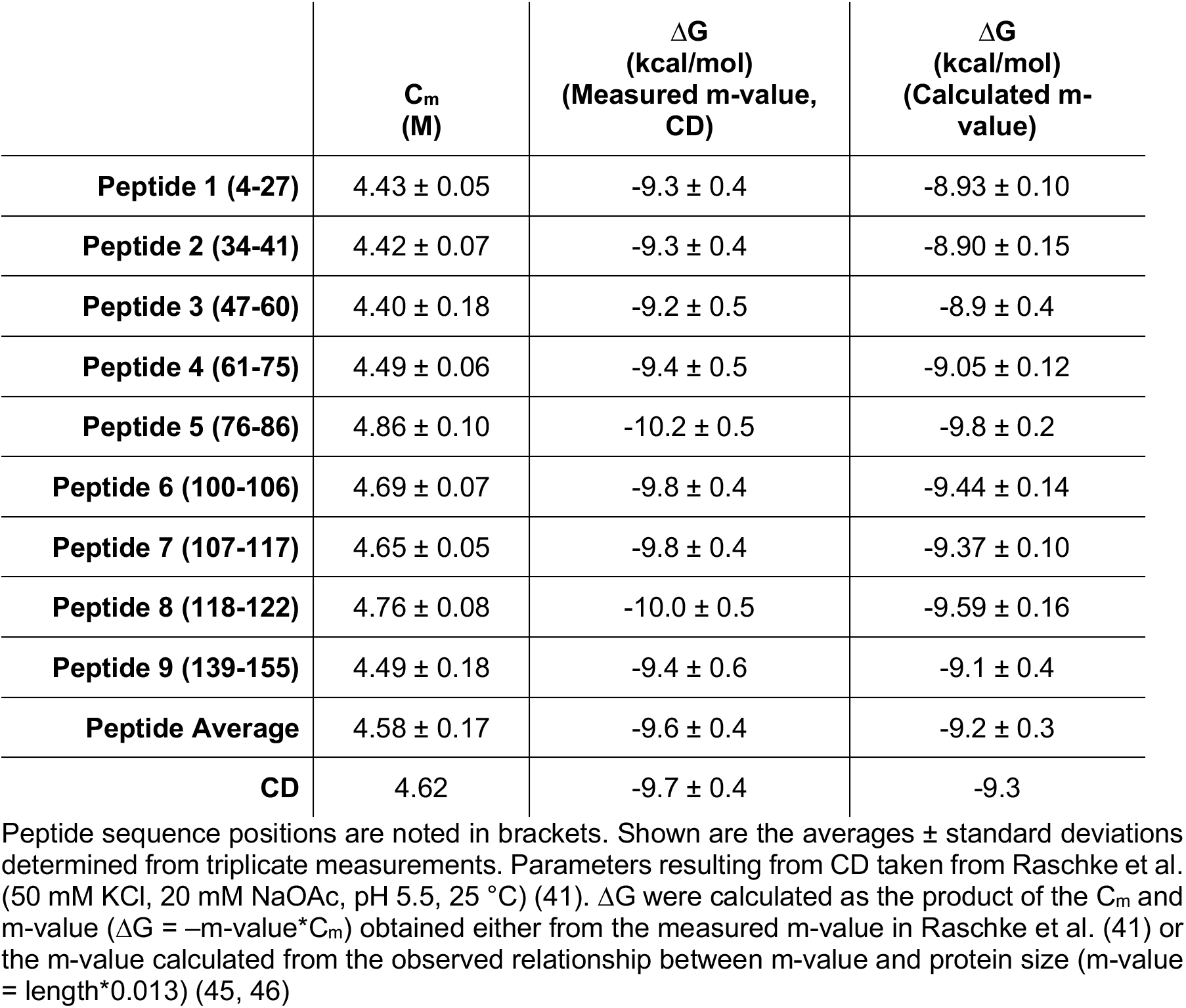
Thermodynamic parameters from ecRNH* XF-MS denaturation monitored at the peptide level.

|  | **C<sub>m</sub><br>(M)** | **∆G<br>(kcal/mol)<br>(Measured m-value,<br>CD)** | **∆G<br>(kcal/mol)<br>(Calculated m-value)** |
|---|---|---|---|
| **Peptide 1 (4-27)** | 4.43 ± 0.05 | -9.3 ± 0.4 | -8.93 ± 0.10 |
| **Peptide 2 (34-41)** | 4.42 ± 0.07 | -9.3 ± 0.4 | -8.90 ± 0.15 |
| **Peptide 3 (47-60)** | 4.40 ± 0.18 | -9.2 ± 0.5 | -8.9 ± 0.4 |
| **Peptide 4 (61-75)** | 4.49 ± 0.06 | -9.4 ± 0.5 | -9.05 ± 0.12 |
| **Peptide 5 (76-86)** | 4.86 ± 0.10 | -10.2 ± 0.5 | -9.8 ± 0.2 |
| **Peptide 6 (100-106)** | 4.69 ± 0.07 | -9.8 ± 0.4 | -9.44 ± 0.14 |
| **Peptide 7 (107-117)** | 4.65 ± 0.05 | -9.8 ± 0.4 | -9.37 ± 0.10 |
| **Peptide 8 (118-122)** | 4.76 ± 0.08 | -10.0 ± 0.5 | -9.59 ± 0.16 |
| **Peptide 9 (139-155)** | 4.49 ± 0.18 | -9.4 ± 0.6 | -9.1 ± 0.4 |
| **Peptide Average** | 4.58 ± 0.17 | -9.6 ± 0.4 | -9.2 ± 0.3 |
| **CD** | 4.62 | -9.7 ± 0.4 | -9.3 |

Peptide sequence positions are noted in brackets. Shown are the averages ± standard deviations determined from triplicate measurements. Parameters resulting from CD taken from Raschke et al. (50 mM KCl, 20 mM NaOAc, pH 5.5, 25 °C) (41). ∆G were calculated as the product of the C<sub>m</sub> and m-value (∆G = –m-value\*C<sub>m</sub>) obtained either from the measured m-value in Raschke et al. (41) or the m-value calculated from the observed relationship between m-value and protein size (m-value = length\*0.013) (45, 46)

**Table 5.**
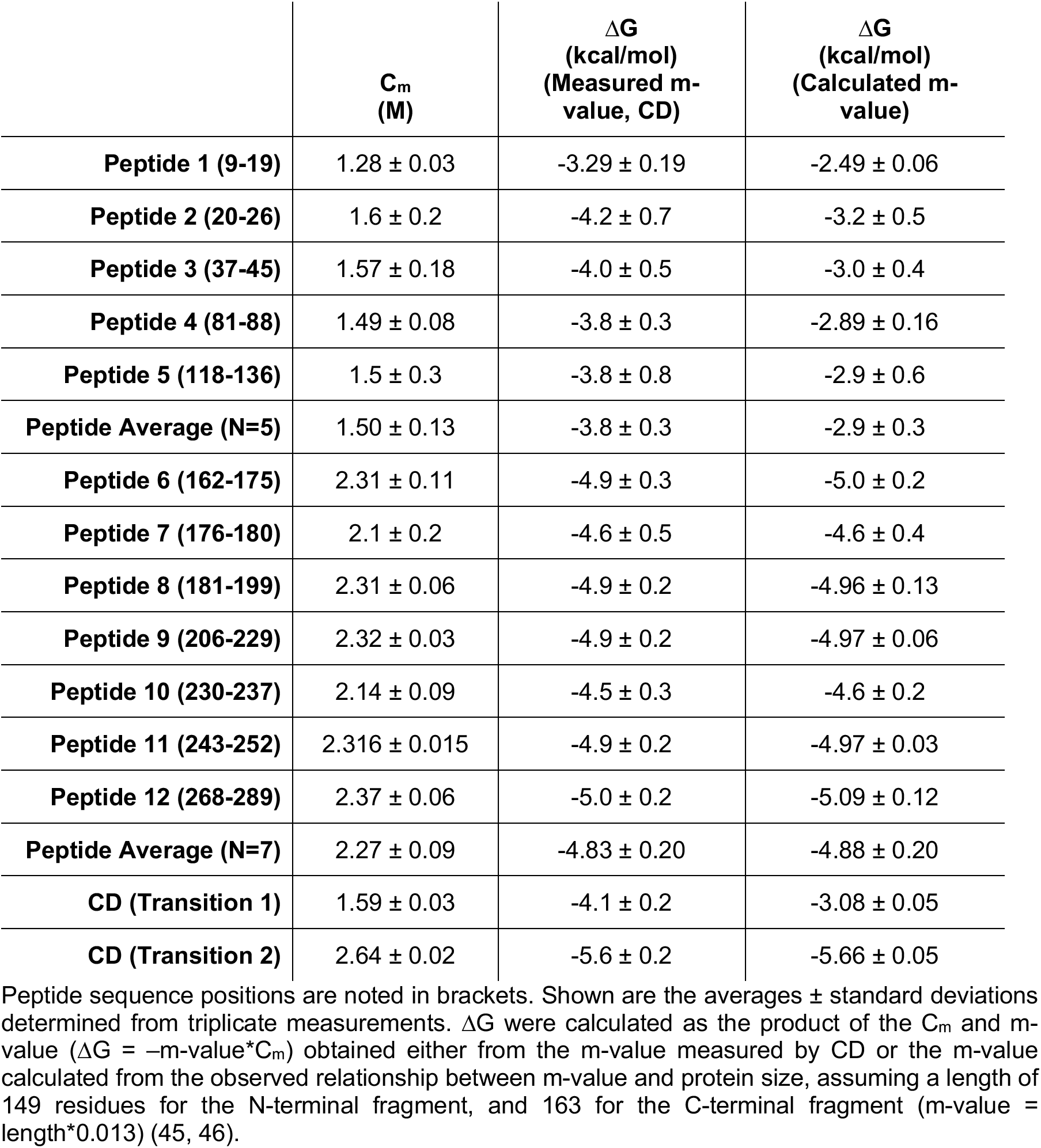
Thermodynamic parameters from SasC 14-17 XF-MS denaturation monitored at the peptide level.

This Cm can be used to determine the 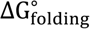 via the m-value (–m-value* C_m_). Using either the m-value predicted from the primary sequence (45, 46), or the known m-value previously measured by CD (41), this modified LEM approach applied to individual peptides returns an average 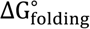 in agreement with the known 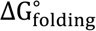 of ecRNH* (Table 4). This result demonstrates the ability of XF-MS, when combined with denaturation, to measure the stability of proteins using local probes (peptides), facilitating the measurement of stability of larger systems in more complex solutions.

The ability to measure stability locally raises the possibility of monitoring non-two-state unfolding. In a perfectly two-state system, the C_m_ should be independent of the observation method (i.e., CD, intact XF-MS, or peptide XF-MS). While as noted above, all peptides report a C_m_ consistent with global unfolding, there appear to be some systematic differences, which may indicate a deviation from two-state behavior. Interestingly, the regions with a lower observed C_m_ are the same regions that are unfolded in both the known kinetic folding intermediate and a known high-energy partially folded equilibrium intermediate of ecRNH* (Fig. 4D) (25, 38, 41, 47, 48). This result demonstrates the ability of XF-MS to monitor protein unfolding using only the change in accessibility of individual peptides, and to potentially detect subtle deviations from two-state behavior.

### The unfolding of individual domains within a multi-domain protein can be monitored with XF-MS

Traditionally, protein denaturant melts are analyzed with a two-state model, which assumes that only the native state and fully unfolded states are appreciably populated under each denaturant condition (19). However, many proteins, especially larger proteins and multi-domain proteins populate partially folded states (22, 23). This can manifest as multiple, clearly resolvable transitions in a denaturant melt for which a two-state model will not fit. However, as evidenced by the above example of a mixture of ecRNH* and T4L*, multiple transitions close to each other can easily be mistaken for a single shallower transition and be adequately, yet inappropriately, fit to a two-state model, resulting in a nonsensical m-value and 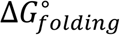. For this reason, it is best practice to test for two-state behavior. This is frequently done by monitoring the unfolding transition using different probes (e.g., comparing CD and Intrinsic Fluorescence). Non-coincident stabilities determined by the different probes is strong evidence of a lack of two-state folding behavior (19).

Peptide XF-MS allows for such a test of two-state behavior, as each peptide denaturant melt provides an independent probe for unfolding. Each peptide reports directly on a specific region, allowing for a structural interpretation. To demonstrate this, we performed denaturant melts monitored by peptide XF-MS using a multi-domain protein, comprised of four sequential domains (domains 14-17) from *S. aureus* surface protein C (SasC) (herein referred to as SasC 14-17) (49, 50). These four domains are predicted to form individual three-helix bundles. When denaturation is monitored by CD, the resulting melt shows two transitions that, while separable, with fewer measurements or additional noise could be mistaken for a broad single transition (Fig. 5A and *SI Appendix*, Fig. S10). Additionally, even with a three-state model, the global signal from CD prevents the structural assignment of the two resulting Cm. When subjected to chemical denaturation monitored by XF-MS, twelve tryptic peptides displayed cooperative transitions (Fig. 5B and *SI Appendix*, Fig. S11). As expected, the observed difference in oxidation between native and unfolded conformations is lower compared to ecRNH*, consistent with the predicted domain structures having smaller hydrophobic cores and therefore a less dramatic change in the solvent-accessible surface area upon unfolding. For SasC 14-17 two classes of peptides are observed, with C_m_ differing by more than 0.6 M urea (Fig. 5B,5D and *SI Appendix*, Fig. S12). When mapped directly onto the predicted structure, the two N-terminal domains show a lower C_m_ than the two C-terminal domains (Fig. 5E). Thus, XF-MS monitored at the peptide level can identify these important cooperativity modules that comprise many proteins and are difficult to detect using probes of global stability.

**Figure 5.**
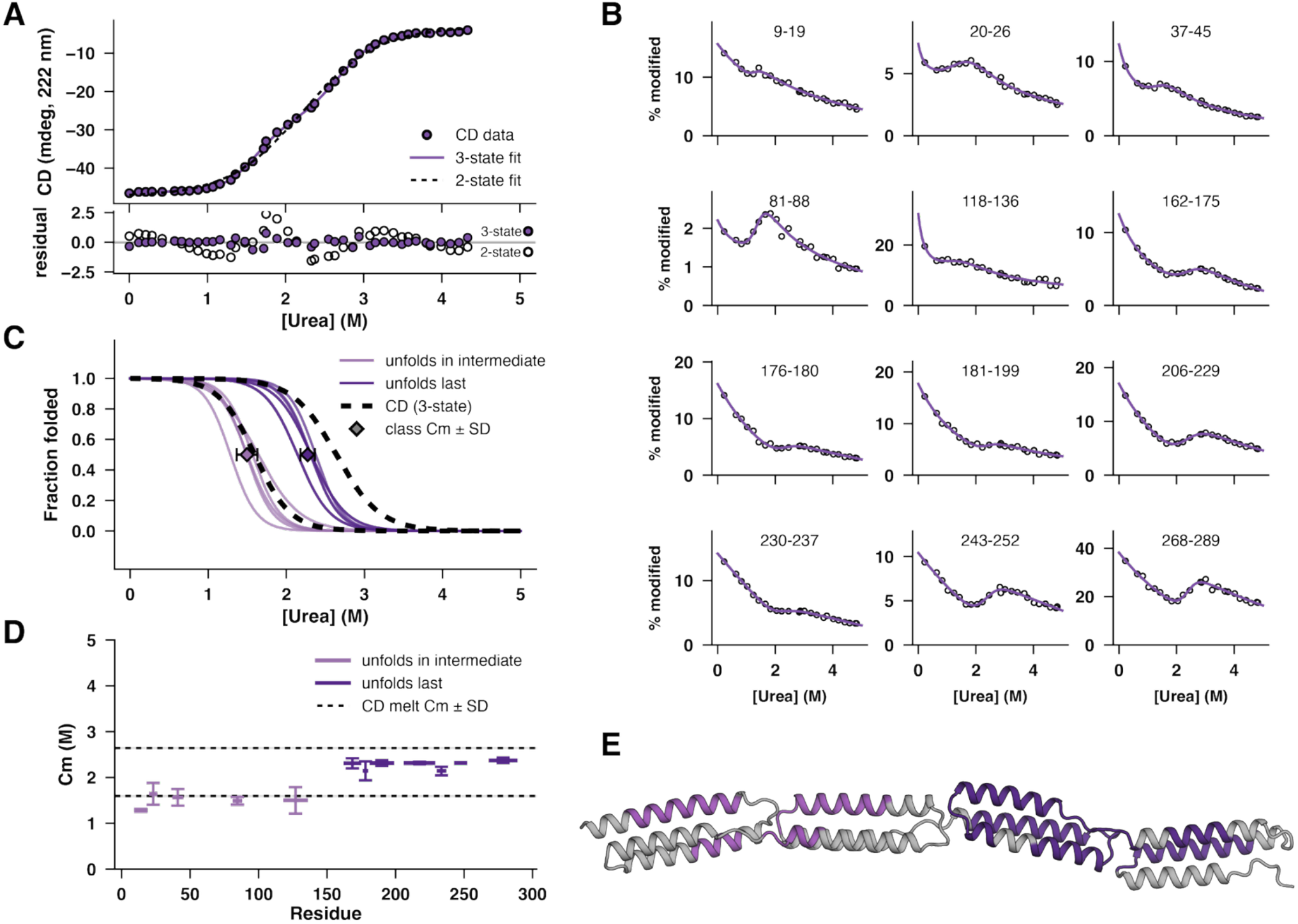
Peptide-level XF-MS reveals multiple cooperative folding units within the multi-domain protein SasC 14-17. (A) A single representative urea denaturation curve (CD 222 nm) of SasC 14-17 (PBS pH7.4, 25 °C). The dotted black line represents a fit to a model assuming two states, folded and unfolded. The solid purple line represents a fit to a model assuming three states, including a partially folded state (see Methods). Shown below are the residuals resulting from the two fits. (B) Percent modified vs. denaturant concentration (PBS pH 7.4, room temperature) for twelve tryptic peptides from SasC 14-17 for a single representative denaturant melt. Residue numbers are indicated in each panel. Purple lines represent fits to a modified LEM (see methods). (C) Overlay of denaturant melts for all tryptic peptides with observable transitions, using the average C_m_ and m-value determined from triplicate measurements for each peptide, normalized to fraction folded. Shown in dotted lines are the expected transitions from a three-state model fit to denaturant melts monitored by CD. (D) Concentration midpoints (C_m_) for each tryptic peptide. Sequence position for each peptide is represented by the x-axis. Error bars represent the standard deviation determined from triplicate measurements. Shown in dotted lines are the expected C_m_ from a three-state model fit to denaturant melts monitored by CD. (E) AlphaFold2 predicted structure of SasC 14-17 (N-terminus on the left). Colored by C_m_ (C_m_ < 2.0 M as light purple, C_m_ > 2.0 M as dark purple, peptides with no cooperative transitions, or regions without peptide coverage, as gray)

## Discussion

In this study, we show that XF-MS combined with chemical denaturation is a powerful method for measuring protein thermodynamic stability and demonstrate XF-MS’s advantages over other (traditional and non-traditional) approaches for monitoring protein denaturation. Specifically, XF-MS requires minimal sample, is applicable to more systems, and has less stringent purity requirements. Compared to traditional methods like circular dichroism (CD) or intrinsic fluorescence, XF-MS requires orders of magnitude less sample to measure stability. Compared to other recently developed methods such as pulse proteolysis (46, 51), SUPREX (52), and SPROX (53, 54), the XF-MS approach described here is amenable to more systems due to the faster labeling kinetics, broader reactivity, and less stringent sample requirements. Importantly, our results also indicate that XF-MS can monitor specific proteins in mixtures, eliminating the need for labor-intensive purification steps and creating new opportunities to study the effects of biological components on protein stability.

In addition to the advantages outlined above, peptide-level XF-MS provides local information that can reveal the detailed energetics of complex protein conformational energy landscapes. Proteins are often assumed to fold completely cooperatively, with only the native state and a globally unfolded state appreciably populated. While this two-state assumption is valid for many proteins <200 residues in length, the median protein is larger, and more likely to populate partially folded states that violate this assumption (55). As demonstrated here, the equilibrium denaturation of these non-two-state proteins can be challenging to monitor, quantify, and structurally describe with traditional spectroscopic approaches. Here, using XF-MS, we observe regional deviations in apparent stability for the single-domain protein ecRNH*, and measure the stability of individual domains within the multi-domain protein SasC 14-17. The ability to measure the apparent stability of multiple individual domains within larger proteins will contribute to more detailed descriptions of inter-domain energetic coupling. These findings underscore the potential of HXF-MS to investigate the stability determinants of larger multi-domain and non-two-state proteins.

The approach outlined in this work is likely to be generalizable and increasingly accessible. Generating a signal difference between folded and unfolded protein in hydroxyl radical footprinting relies on buried hydrophobic residues in the native state, a nearly universal quality of globular proteins. This feature makes it more broadly applicable compared to other MS-based methods that rely instead on the selective aggregation of the unfolded state, like Thermal Proteome Profiling (TPP), or changes in protease susceptibility, like Limited Proteolysis (LiP) (56–63). Additionally, unlike TPP and LiP, XF-MS combined with chemical denaturation provides rigorous thermodynamic information. TPP, LiP, and other similar methods probe many proteins in parallel by mass spectrometry. XF-MS can similarly take advantage of advances made by the proteomics field, further expanding the range of systems for which quantitative thermodynamic parameters can be measured.

Despite its unique capabilities, the need for specialized equipment limits the widespread adoption of XF-MS for measurements of protein stability. However, mass spectrometers are increasingly available in core facilities, bench-top hydrogen peroxide-based hydroxyl radical footprinting instruments have long been in use to great effect, and plasma-based instruments are now commercially available (36, 64). These alternative sources of radicals present their own strengths and weaknesses, but in general, require even shorter labeling times and have been effectively used to study proteins *in vivo* (65–67). The accessibility of these instruments will facilitate the adoption of hydroxyl radical footprinting for a broader range of systems in a broader range of research settings.

It is worth noting that measuring protein thermodynamic stability doesn’t always require chemical or temperature denaturation. Amide hydrogen exchange has successfully measured protection at the level of individual residues, which is often a proxy of stability (24, 25, 68). Similarly, proteolysis under native conditions can report on the populations of high-energy states (26, 69). Recent work combining yeast display or cDNA display with proteolysis to estimate protein stability has resulted in a high-throughput exploration of protein stability sequence space for small single-domain proteins (13, 70, 71). These native-state approaches have advantages over denaturation-based approaches. Primarily, they require no extrapolation from non-native conditions and therefore do not require making the same two-state assumptions that are often unknowingly violated in chemically induced denaturation. Native-state methods also report on conformational heterogeneity beyond global unfolding, including local unfolding and—especially in the case of amide hydrogen exchange—smaller scale conformational fluctuations. However, these approaches also have their limitations. Knowing whether a given native-state measurement is reporting on global unfolding, local unfolding, or smaller scale fluctuations is not always clear. Additionally, not all native-state methods are applicable to all systems. Native-state proteolysis, for example, is limited to proteins without cleavable regions in their native-state ensemble. Ultimately, denaturation and native-state approaches for measuring conformational heterogeneity are both useful and the choice of which method is most appropriate or informative largely depends on the idiosyncrasies of each protein of interest and the questions being asked.

Combining XF-MS and chemical denaturation is a powerful tool for studying protein thermodynamic stability in complex biological systems. Its advantages over traditional methods and compatibility with a wide range of systems make it a valuable addition to the toolbox of techniques for investigating protein structure, stability, and function. Future work will optimize this method for high-throughput applications, expand the range of proteins and biological systems that can be studied, and elucidate the molecular mechanisms underlying protein stability and folding dynamics in native environments.

## Materials and Methods

### Protein purification

For the purification of ecRNH*, the clarified lysate was purified first over a HiTrap Heparin column (GE Healthcare) at pH 8. Peak fractions were pooled and diluted 3-fold with doubly deionized water. The pH of the solution was then adjusted to 5.5 using dilute HCl. The resulting aggregates were removed by another round of centrifugation and filtration. The filtered sample was further purified over a HiTrap S column (GE Healthcare). Protein was then concentrated and buffer-exchanged into ammonium bicarbonate for subsequent freeze-drying and storage. Prior to use, the freeze-dried protein was resuspended in 20 mM NaOAc, 50 mM KCl, pH 5.5, and filtered (0.22 um).

For the purification of T4L*, clarified lysate was loaded onto a Capto S column, washed with 25 mM Tris, 10 mM NaCl, pH 8.0, and eluted with a gradient from 10 mM to 300 mM NaCl (25 mM Tris pH 8.0). Fractions containing T4L* were pooled, concentrated, and run on a S200 (GE Healthcare) column in PBS pH 7.4 (Sigma Aldrich P4417). Protein was stored at 4 °C until use.

For the purification of His6-Prescission-SasC(14–17), clarified lysate was loaded onto a Ni-NTA, washed with 20 mM Tris, 500 mM NaCl, pH 8.0, and eluted with a gradient from 0 mM to 500 mM imidazole (20 mM Tris, 500 mM NaCl, pH 8.0). Fractions containing His6-Prescission-SasC(14–17) were pooled and dialyzed into 25 mM Tris, 150 mM NaCl, 0.5 mM TCEP, pH 8.0. Prescission protease was added and equilibrated overnight at 4 °C, followed by an additional Ni-NTA column and collection of the flowthrough. Fractions containing SasC(14–17) were pooled, dialyzed into 25 mM Tris, pH 8.0, then loaded onto a Q-Xl column, washed with 25 mM Tris pH 8.0, and eluted with a gradient from 0 mM to 1000 mM NaCl (25 mM Tris pH 8.0). Fractions containing SasC(14–17) were pooled concentrated, and run on a S200 (GE Healthcare) column in PBS pH 7.4 (Sigma Aldrich P4417). Protein was stored at 4 °C until use.

### Preparation of soluble E. coli lysate

E. coli K12 was grown overnight at 37 °C in 50 ml of LB media and harvested by centrifugation. The cell pellet was resuspended in 50 ml of 20 mM Tris–HCl, 250 mM NaCl, 10 mM EDTA pH 8.0 then pelleted again by centrifugation. The washed cell pellet was resuspended in 10 ml of 20 mM Tris–HCl, 50 mM NaCl, 1 mM EDTA pH 8.0 with added protease inhibitor and lysed by lysozyme treatment and repeated freeze-thaw cycles. Cell debris was then pelleted (ss34, 14k rpm, 30 minutes at 4 °C) and the resulting supernatant was filtered (0.22 um). Nucleic acids were digested by incubating supernatant with 0.1 mg/ml DNase I and 0.1 mg/ml RNase A. MgCl_2_ and CaCl_2_ were added to 2.5 mM and 1.0 mM, respectively, for this digestion reaction. To remove small metabolites and digested nucleic acids, the lysate was buffer exchanged using an amicon concentrator with a 3 kDa cutoff, first against 20 mM Tris–HCl, 250 mM NaCl, pH 8.0 and then against 20 mM NaOAc, 50 mM KCl, pH 5.5. The lysate was sterilized by passing through a 0.22 μm syringe filter and stored at −20 °C until used.

### **X-**ray hydroxyl radical footprinting

XF experiments were carried out at the Advanced Light Source beamline 3.3.1 using a standard microfluidic set-up (28, 43, 72). Protein concentrations were as follows: 5 µM for all isolated proteins, 2.5 µM each for the combined ecRNH* and T4L* experiments, and 1 µM ecRNH* in lysate. ecRNH*, Lysate, and T4L* experiments were in 20 mM NaOAc, 50 mM KCl, pH 5.5. SasC experiments were in PBS (Sigma Aldrich P4417). Samples were subject to 500 or 1000 µs of X-ray exposure then immediately quenched with methionine amide (final concentration of 5 mM) to prevent secondary oxidations and stored at -80 °C until LC-MS sample preparation and analysis.

### Intact protein mass spectrometry and analysis

Intact proteins were injected directly at a flow rate of 50 uL/min in 95% Buffer A (0.1% Formic Acid) 5% buffer B (100% Acetonitrile, 0.1% Formic Acid) onto a mAbPac column (Thermo Fisher) stored in a column oven maintained at 50 °C. Residual buffer, urea, and salts were removed by maintaining flow for 4 minutes with a valve diverting to waste. Proteins were then eluted with a gradient of 5-20% buffer B at a flow rate of 50 μL/min over 30 seconds, 20-70% buffer B over 4 minutes, then 70-90% over 30 seconds. The column was then subjected to a sawtooth wash and equilibrated at 5% buffer B prior to the next injection. Proteins were eluted directly into a Q Exactive Orbitrap Mass Spectrometer operating in positive mode (HESI source settings: Spray voltage 4.0 kV, Capillary temp. 325 C, S-lens RF level 70, Aux gas heater temp. 150 °C, In-source CID 15.0 eV — MS Settings: 6 microscans, resolution 140000, AGC target 5e6, maximum IT 100 ms, scan range 600-3000 m/z).

Intact mass spectra were summed across the entire chromatographic peak, then exported from Xcalibur and imported into Unidec (73, 74). Spectra were deconvoluted with Unidec, to remove charge state redundancy and reduce noise. Peak width was manually adjusted to match the isotopically resolved data. Spectra were sampled at every 0.1 Da. Charge state distributions were automatically smoothed. Deconvoluted spectra were quantified by calculating the average mass across a mass region spanning all observed modifications for a given intact protein.

### Peptide mass spectrometry and modification analysis

Prior to digestion, samples were diluted 5-fold with 100 mM Tris pH 8.0. Trypsin (Promega) was added at a protease:protein ratio of 1:100, and samples were incubated overnight at 37 °C. Digestion was quenched with the addition of formic acid to a final concentration of 1%. Following digestion, peptides were loaded at a flow rate of 50 uL/min in 99% Buffer A (0.1% Formic Acid) 1% buffer B (100% Acetonitrile, 0.1% Formic Acid) onto a Pepmap C18 column (Thermo Fisher 164711) stored in a column oven maintained at 50 °C. Residual buffer, urea, and salts were removed by maintaining flow for 4 minutes with a valve diverting to waste. Peptides were then eluted with a gradient of 1-40% buffer B at a flow rate of 50 μL/min over 15 minutes, and then of 40-90% buffer B over 30 seconds. The column was then subjected to a sawtooth wash and equilibrated at 1% buffer B prior to the next injection. Peptides were eluted directly into a Q Exactive Orbitrap Mass Spectrometer operating in positive mode (HESI source settings: Spray voltage 3.5 kV, Capillary temp. 150 C, S-lens RF level 70, Aux gas heater temp. 150 °C — MS Settings: resolution 70000, AGC target 3e6, maximum IT 50 ms, scan range 150-2000 m/z — dd-MS^2^ settings: resolution 17500, AGC target 1e5, maximum IT 50 ms, loop count 10, isolation window 2.6 m/z, NCE 28, charge state 1 and ≥7 excluded, dynamic exclusion of 2 seconds). LC and MS methods were run using Xcalibur 4.1 (Thermo Scientific)

Modified and unmodified peptide identification and quantification were performed using the Byos Oxidative Footprinting platform (Protein Metrics Incorporated) incorporating both the Byonic search engine and Byologic quantification program. All peptide experiments with fragmentation data were searched for tryptic peptides originating from the search library. The search library consisted of a fasta file containing sequences for all proteins present in solution. Half-tryptic peptides and peptides with missed tryptic cleavages were present, but with lower intensity, and were excluded from further analysis. For a given tryptic peptide, all detected modifications were quantified if observed in any sample. Each modification type (i.e. +16 Da, +32 Da, +48 Da, etc.) was separately quantified by summing the area of all chromatographic peaks with that mass, regardless of whether an MSMS identification was made at that retention time. Chromatographic peaks with isotopic distribution that did not match the unmodified peak, or for which MSMS had identified as a different peptide, were not quantified. The final percent modified was the sum of all modified peak areas divided by the sum of all modified and unmodified peak areas.

### Circular dichroism

Circular dichroism experiments were performed with either an Aviv 410 spectrometer (ecRNH* and T4L*) or a Jasco J-1500 (SasC 14-17). Each sample contained 40 μg/mL protein (either ecRNH*, T4L*, an equimolar mixture of both, or SasC 14-17) with varying concentrations of urea. CD melts for ecRNH* and T4L* were performed in 20 mM NaOAc 50 mM KCl pH 5.5. CD melts for SasC were performed in PBS, pH 7.4. The signal at 222 nm was measured in a 1-cm path length Starna Cells cuvette at 25°C with stirring, averaged over one minute.

### Extraction of thermodynamic parameters from denaturation melts

Denaturant melts, for both circular dichroism and XF-MS, were analyzed using the linear extrapolation model, resulting in both a C_m_ and m-value describing the unfolding transition and parameters describing the native and denatured baselines. For data with linear baselines (CD, intact XF-MS), the following equation was used:

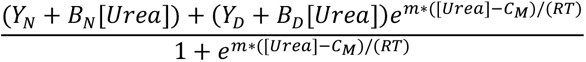

Where m and C_m_ are the m-value and concentration midpoint, Y_D_ and Y_N_ represent the denatured and native baseline Y-intercepts, B_D_ and B_N_ represent the denatured and native baseline slopes, and where R is the gas constant and T is the temperature in Kelvin. R and T were fixed at 1.987E-3 kcal/(mol*K) and 293 K, respectively.

For denaturant melts of SasC 14-17 by CD, with two transitions, the following equations were used:

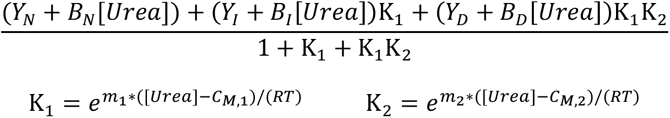

For peptide XF-MS melts with non-linear baselines, the following equation was used:

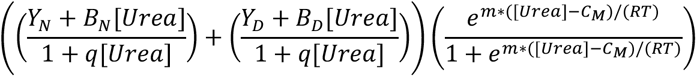

With the same definitions as above and where q represents a peptide-specific urea quenching term.

Parameters were determined by non-linear least squares fitting of the data to the appropriate model with scipy.optimize in Python 3.

## Acknowledgments

We thank the entire Marqusee lab for experimental advice and support. This work was supported by funding from the NIH (S.M.) and the NSF (S.M.). S.M. is a Chan-Zuckerberg Biohub investigator.

