## Supporting Information for "Determination of both global and local protein stability in complex environments using X-ray hydroxyl radical footprinting"

**This PDF file includes:**

Figures S1 to S12

### Figures

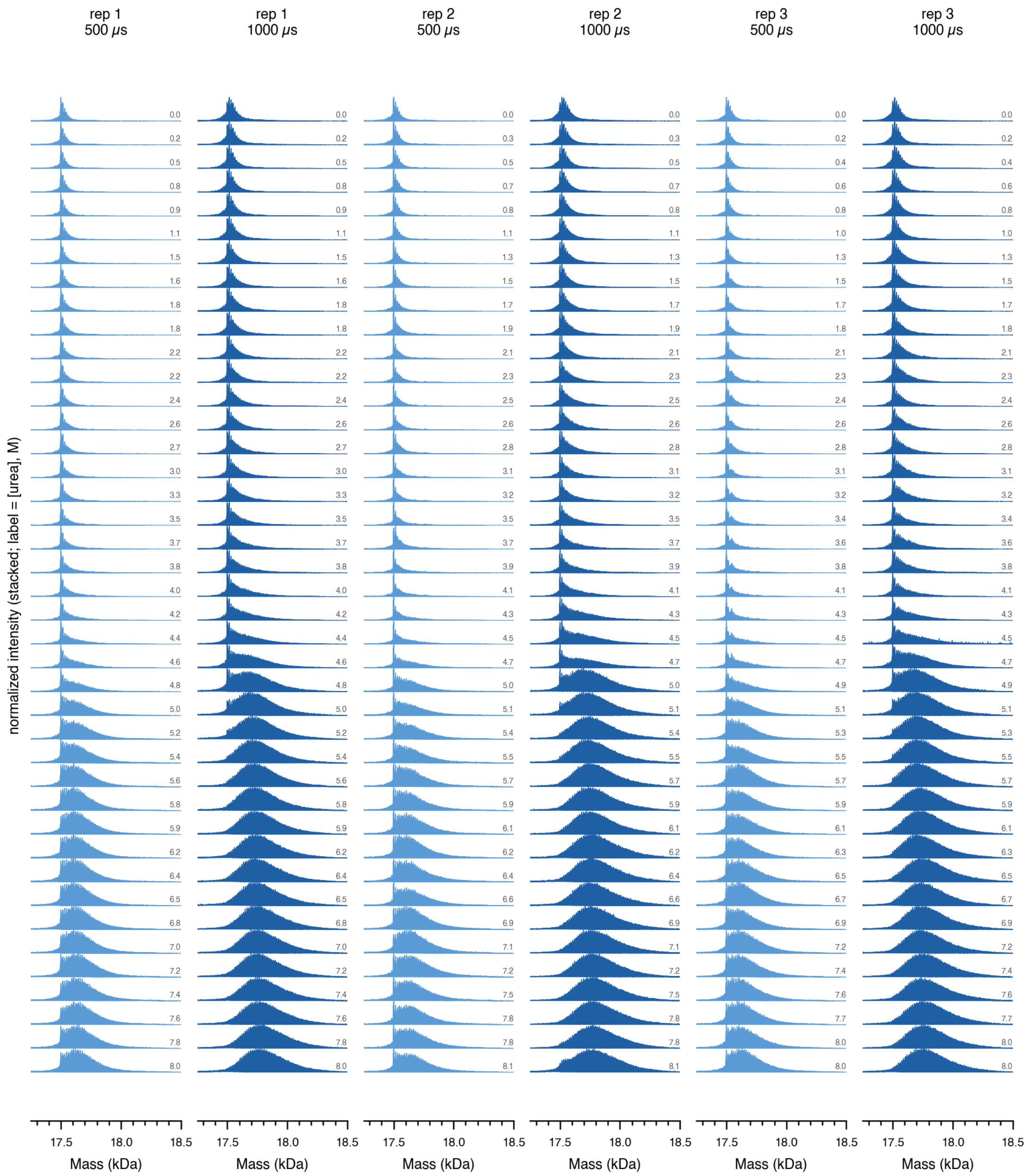


**Fig. S1. Denaturant-dependent oxidation of ecRNH* is reproducible across replicates and exposure times.** Deconvoluted intact-mass spectra of ecRNH* (50 mM KCl, 20 mM NaOAc, pH 5.5, room temperature) are shown for three replicates at 500 and 1000 μs exposure across the full urea series, with molar urea concentrations indicated to the right of each spectrum. The progressive increase in mass reflects increased oxidation of side chains that become exposed upon unfolding.

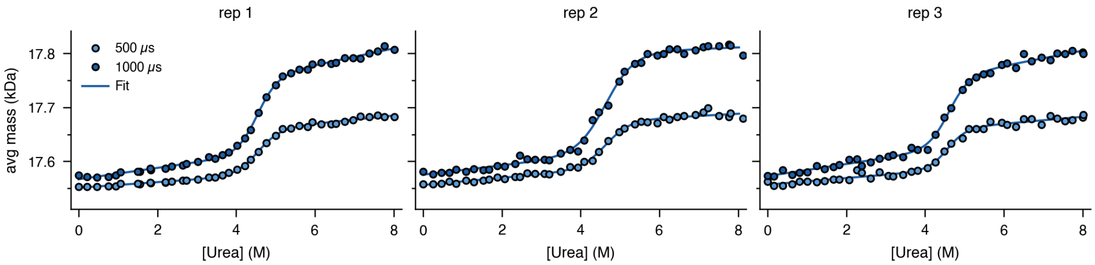


**Fig. S2. Intact-mass denaturant melts of ecRNH* are reproducible across replicates and exposure times.** The average deconvoluted mass of ecRNH* (50 mM KCl, 20 mM NaOAc, pH 5.5, room temperature) is plotted versus urea for three replicates at 500 µs (light) and 1000 µs (dark), with two-state linear-extrapolation-model (LEM) fits (solid lines). The main-text Fig. 1B shows a single representative replicate; fitted parameters for all replicates and pulse lengths are reported in Table 1.

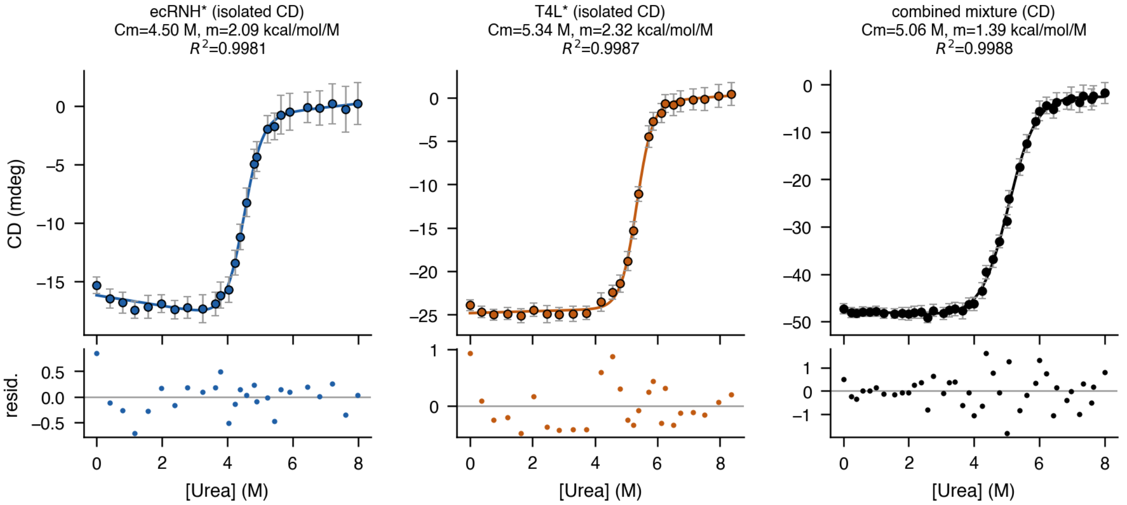


**Fig. S3. CD cannot resolve the two components of an ecRNH*/T4L* mixture and returns a nonphysical stability.** CD at 222 nm (mdeg) is plotted versus urea for isolated ecRNH* (blue), isolated T4L* (orange), and the 1:1 ecRNH*/T4L* mixture (black), each fit to a two-state LEM (solid line) with residuals shown below and the fitted Cm, m-value, and $R^2$ above each panel. The mixture is adequately fit as a single two-state transition yet returns a ΔG°folding of −7.0 kcal/mol, inconsistent with either component (Table 2).

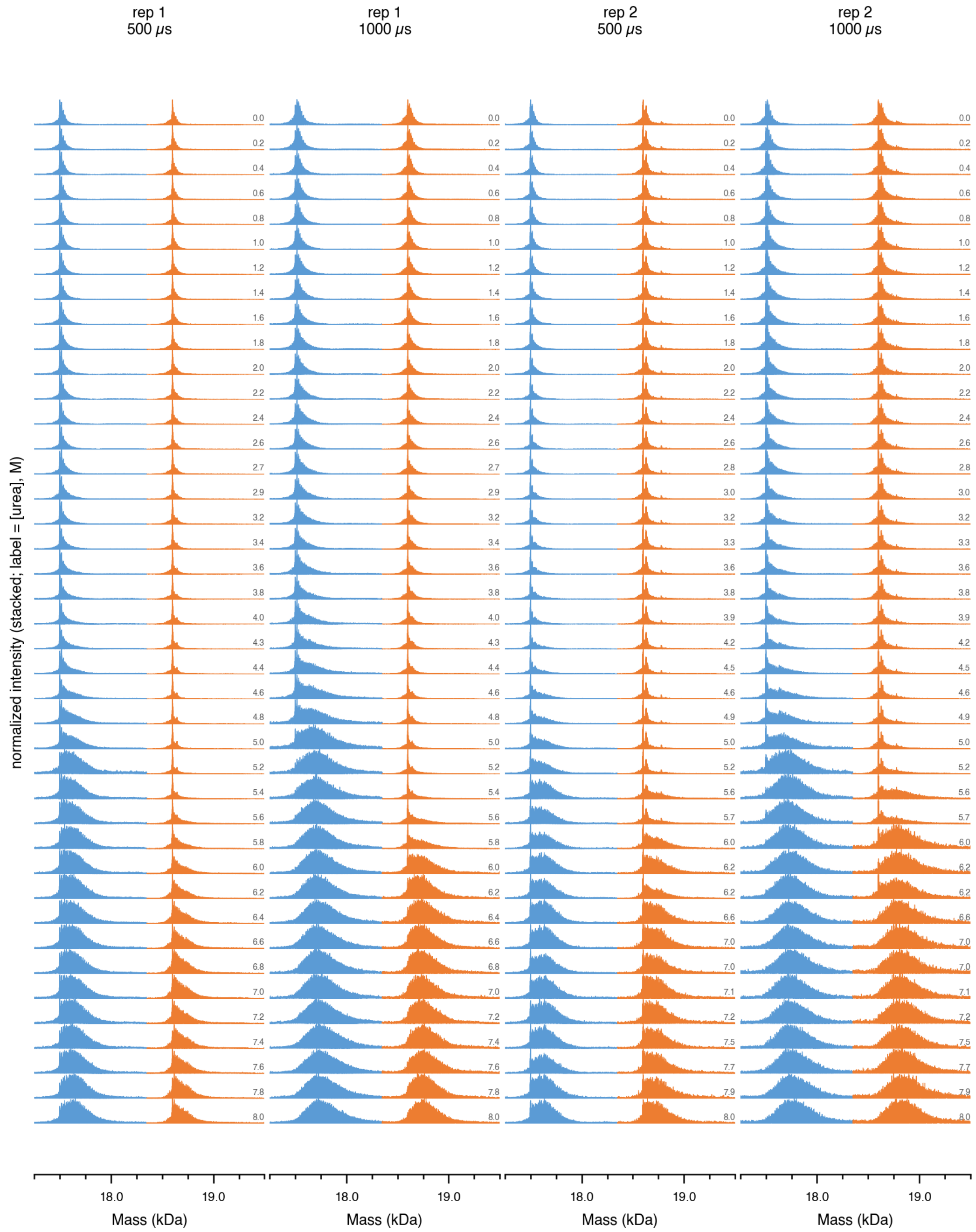


**Fig. S4. ecRNH* and T4L* occupy non-overlapping mass regions throughout the mixture denaturant series.** Deconvoluted intact-mass spectra of the 1:1 mixture (50 mM KCl, 20 mM NaOAc, pH 5.5, room temperature) are shown for two replicates at 500 and 1000 μs exposure across the full urea series, with molar urea concentrations indicated to the right of each spectrum. The ecRNH* (blue) and T4L* (orange) mass regions are normalized separately, and each shifts at a distinct denaturant concentration.

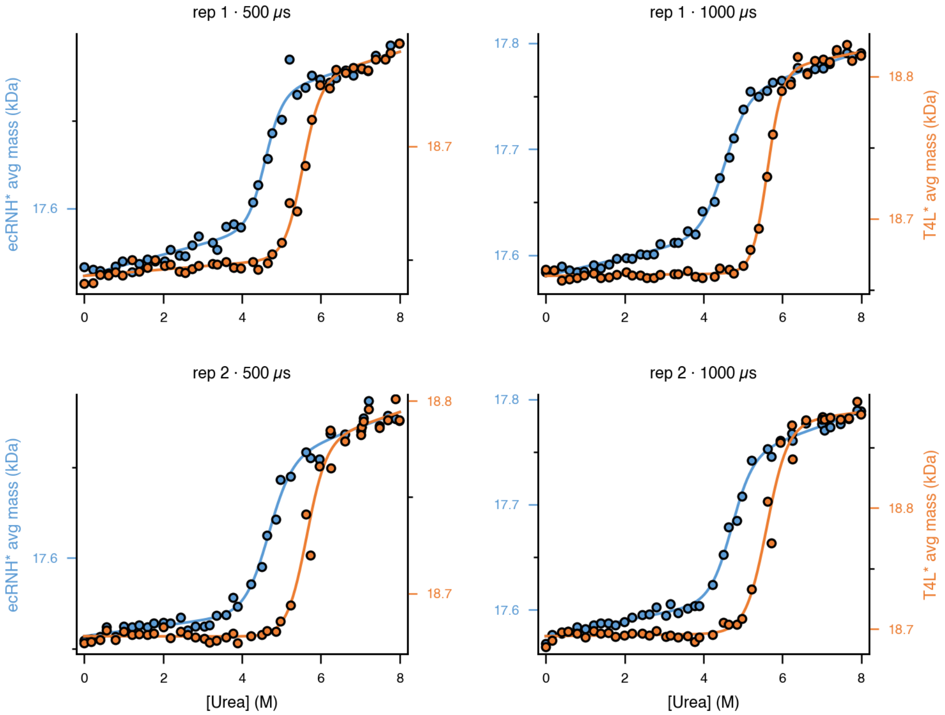


**Fig. S5. XF-MS independently resolves the unfolding of ecRNH* and T4L* in a mixture across all replicates.** The average deconvoluted mass of ecRNH* (blue, left axis) and T4L* (orange, right axis) from the 1:1 mixture is plotted versus urea for each replicate and exposure time (500 and 1000 μs), with two-state LEM fits (solid lines). The resulting stabilities match the isolated proteins (Table 2); the main-text Fig. 2B shows a single representative melt.

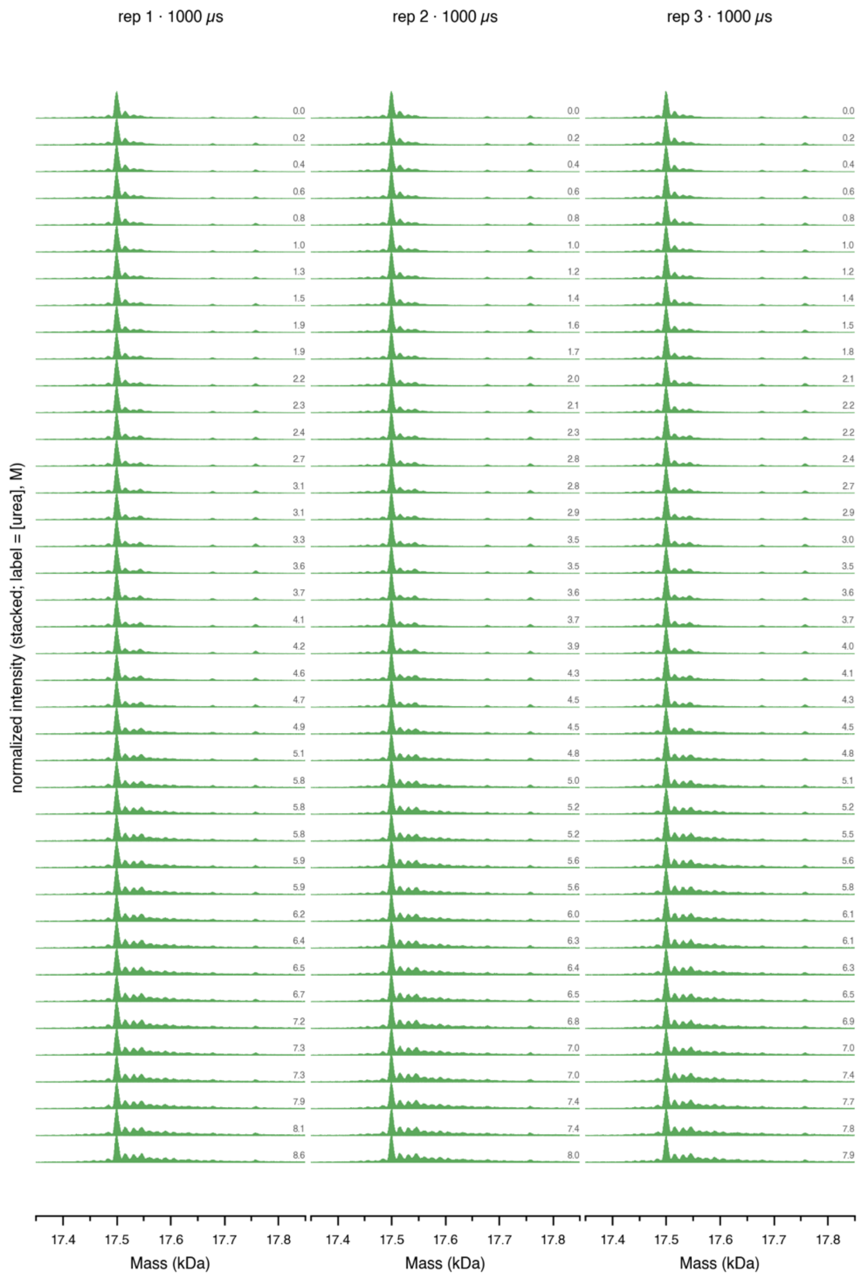


**Fig. S6. ecRNH* oxidation in the presence of clarified lysate tracks denaturant but is attenuated by radical quenching across all replicates.** Deconvoluted intact-mass spectra of 1 μM ecRNH* in the presence of clarified *E. coli* lysate (50 mM KCl, 20 mM NaOAc, pH 5.5, room temperature) are shown for three replicates at 1000 μs exposure across the full urea series, with molar urea concentrations indicated to the right of each spectrum. The mass shift is markedly smaller than for isolated ecRNH* (Fig. S1), reflecting radical quenching by lysate components.

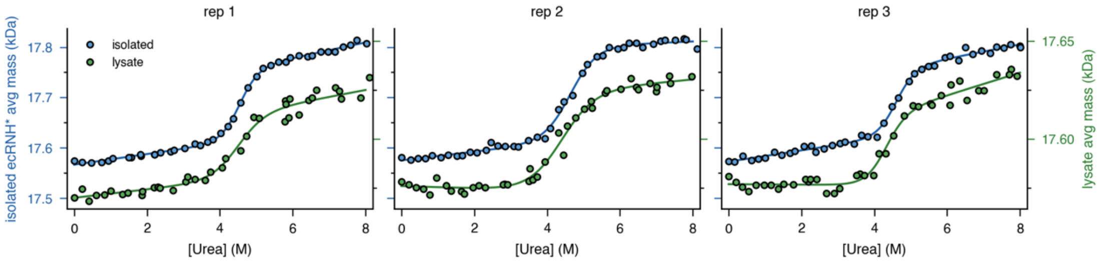


**Fig. S7. XF-MS resolves the ecRNH* denaturant melt in the presence of clarified lysate across replicates despite attenuated signal.** The average deconvoluted mass of isolated ecRNH* (blue, left axis) and ecRNH* in the presence of clarified *E. coli* lysate (green, right axis) is plotted versus urea for three replicates (1000 μs), with two-state LEM fits (solid lines); note the compressed right axis. Fitted parameters are reported in Table 3.

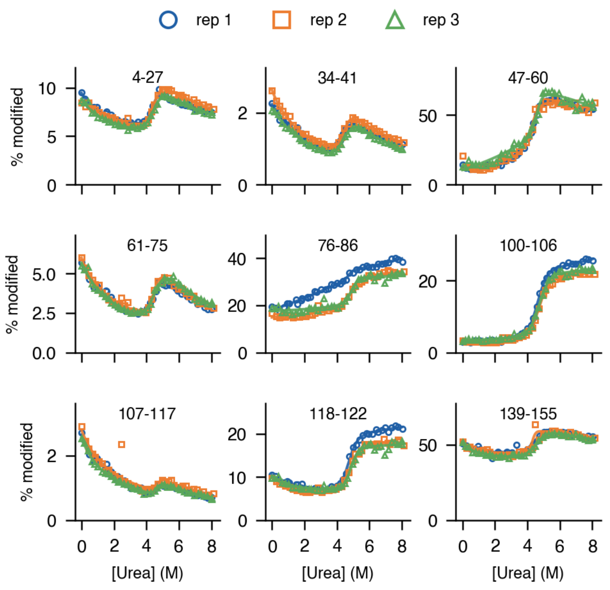


**Fig. S8. Peptide-level denaturant melts of ecRNH* are reproducible across replicates.** Percent modification (50 mM KCl, 20 mM NaOAc, pH 5.5, room temperature, 1000 μs exposure) is plotted versus urea for the nine ecRNH* tryptic peptides with cooperative transitions, labeled by residue range, with replicates overlaid (rep 1, blue circles; rep 2, orange squares; rep 3, green triangles) and fits to a modified LEM (see Methods). The main-text Fig. 4A shows a single representative replicate.

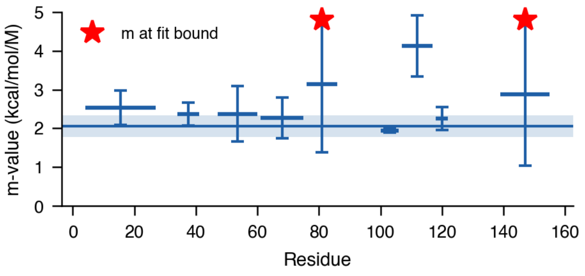


**Fig. S9. Peptide m-values for ecRNH* agree with the global value but are individually poorly determined.** Each point is the mean ± SD across replicates plotted at the peptide midpoint residue; the horizontal line and shaded band give the reference m-value ± SD from the intact/CD measurement. Red stars mark peptides whose m-value reached the fit bound, reflecting the added uncertainty of the seven-parameter modified LEM.

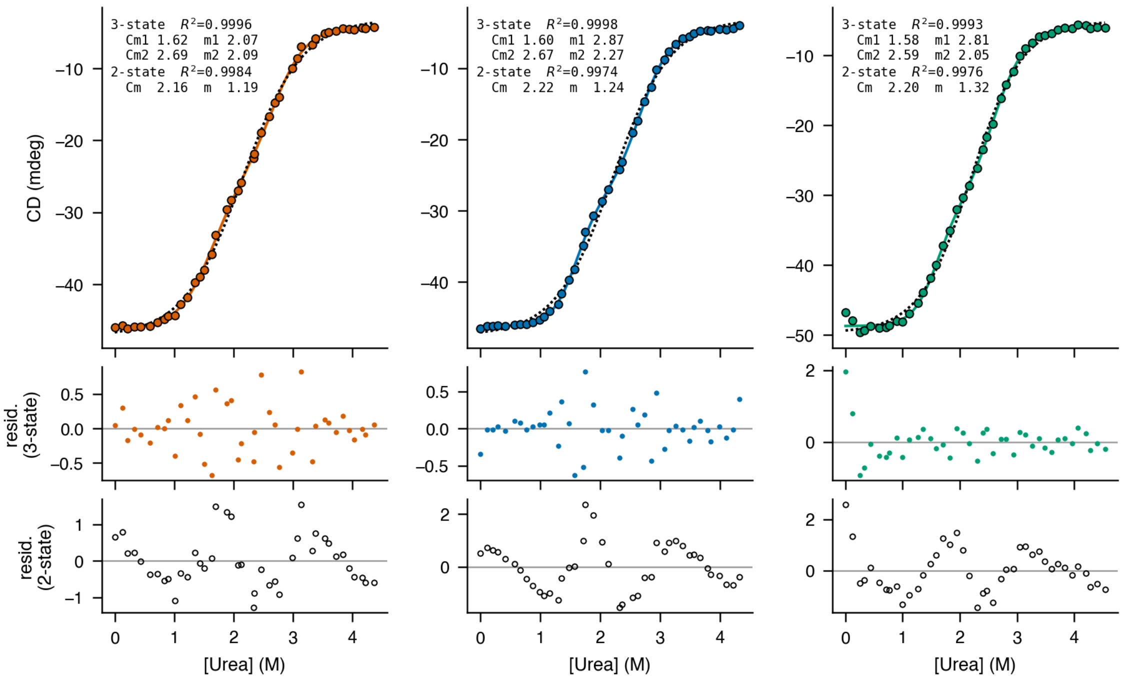


**Fig. S10. A three-state model best describes the SasC 14-17 CD denaturant melts.** Three equilibrated SasC 14-17 CD melts (PBS, pH 7.4, 25 °C) are each fit independently to a two-state (dotted black) and a three-state (solid) LEM; CD at 222 nm (mdeg) is plotted versus urea, the fitted Cm/m-values and $R^2$ for both models are listed in each panel, and the three-state then two-state residuals are stacked below. The systematic structure in the two-state residuals, absent from the three-state residuals, confirms non-two-state unfolding with midpoints near Cm1 ≈ 1.6 M and Cm2 ≈ 2.6 M.

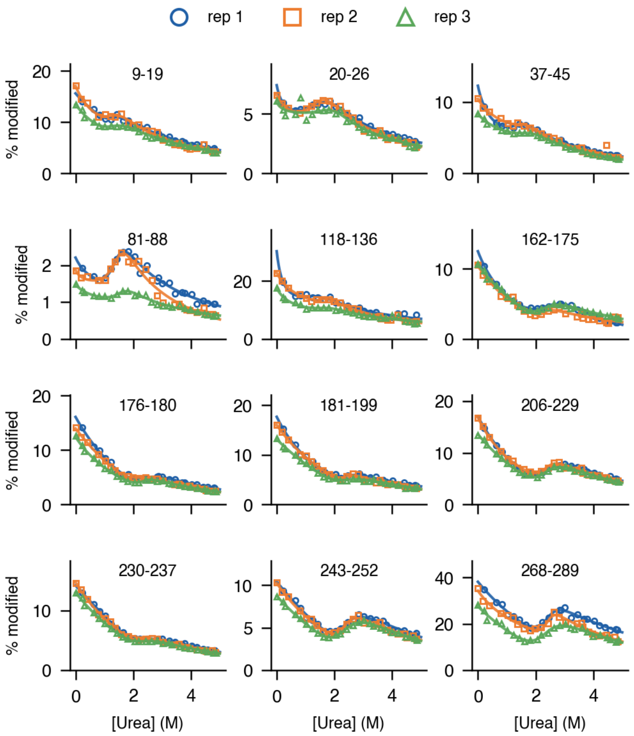


**Fig. S11. Peptide-level denaturant melts of SasC 14-17 are reproducible across replicates and fall into two Cm classes.** Percent modification (PBS, pH 7.4, room temperature, 1000 μs exposure) is plotted versus urea for the twelve SasC 14-17 tryptic peptides with cooperative transitions, labeled by residue range, with replicates overlaid (rep 1, blue circles; rep 2, orange squares; rep 3, green triangles) and fits to a modified LEM (see Methods). The main-text Fig. 5B shows a single representative replicate.

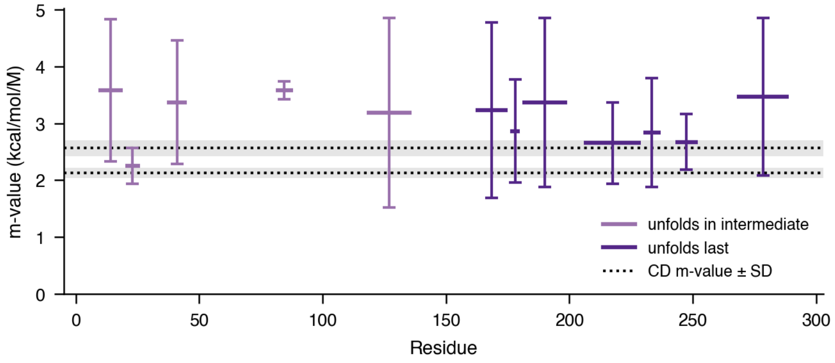


**Fig. S12. Fitted m-values for the SasC 14-17 peptides.** Each point is the mean ± SD across replicates plotted at the peptide midpoint residue, colored by class: peptides that unfold in the intermediate transition (lower Cm, light purple) and those that unfold last (higher Cm, dark purple). Dotted lines mark the two pooled CD m-values, each with a shaded ± SD band.
